# TGIF is a Golgin-like protein required for Golgi structural maintenance and function in *Toxoplasma gondii*

**DOI:** 10.64898/2026.05.21.726867

**Authors:** Camille Pearce, Aoife T. Heaslip

## Abstract

The Golgi is an essential organelle that serves as a central hub for endomembrane trafficking. In the protozoan parasite *Toxoplasma gondii*, a single Golgi stack is essential for parasite survival; however, the molecular determinants governing Golgi structure and function remain poorly understood. Here, we characterize a Golgi-associated protein that is required for Golgi integrity and function, which we named Toxoplasma Golgi Integrity Factor (TGIF). Loss of TGIF disrupts parasite replication and natural egress and is lethal to the parasite. To investigate the impact of TGIF depletion on secretory protein trafficking, we adapted a fluorescence-based pulse–chase assay to monitor the synthesis and trafficking of microneme and rhoptry proteins. We found that loss of TGIF significantly impaired the synthesis and trafficking of microneme and rhoptry neck proteins, whereas trafficking of rhoptry bulb proteins was minimally affected. These findings suggest that rhoptry bulb proteins may traffic independently of canonical Golgi-dependent pathways. Collectively, our study provides new insight into the mechanisms of Golgi-mediated trafficking in *T. gondii* and identifies TGIF as a critical regulator of parasite secretory pathway organization and function.

## Introduction

*Toxoplasma gondii* is an obligate intracellular parasite and the causative agent of Toxoplasmosis. Parasite growth and pathogenesis are dependent on completion of the lytic cycle where parasites attach to and invade host cells and replicate intracellularly within a specialized organelle called the parasitophorous vacuole (PV). Parasite egress results in destruction of the infected cell^1^. Completion of this lytic cycle depends on protein trafficking through the endomembrane system. First, three secretory organelles, the micronemes, rhoptries, and dense granules that contain distinct sets of secreted cargo proteins, are required for invasion, egress, organization of PV and host cell manipulation^2^. Additionally, during parasite division, proteins trafficked through the endomembrane pathway contribute material to the formation inner membrane complex (IMC), a cytoskeletal structure consisting of flattened membranous sacs. Thus, processing and trafficking of cargos from their site of synthesis in the ER through the Golgi apparatus to these subcellular compartments is crucial for *T. gondii* survival.

The Golgi apparatus is an essential organelle present in all eukaryotes and is the central hub of protein trafficking in the endomembrane system. In most species, the Golgi is composed of flattened sac-like plates called cisternae that form a stack ^3–6^. The *cis* face of the Golgi receives cargo proteins from the endoplasmic reticulum (ER). As cargo molecules transit sequentially through the Golgi cisternae, they undergo post-translational modifications mediated by Golgi-resident enzymes. These modifications, including glycosylation, phosphorylation, and proteolysis, are essential for proper protein folding, function, and subcellular localization^4,6–10^. At the *trans* side of the Golgi cargo is sorted and packed into vesicles for trafficking to other subcellular compartments, and for secretion^7^. Retrograde trafficking of resident enzymes is critical for maintaining cisternal identity, and the proper distribution of Golgi-resident enzymes^11^.

Protein trafficking between Golgi cisternae is mediated by transport vesicles that are abundant around the Golgi stack^12^. Vesicle budding at Golgi cisternae is controlled by the COPI coatomer complex. Once formed, several classes of Golgi tethering complexes function to guide incoming vesicles to target membranes. In metazoans, golgins are long coiled-coil domain containing tethers that are anchored at the cisternae and can extend hundreds of nanometers into the cytosol depending on their length^13–15^. Anchoring golgins to the Golgi membranes occurs through transmembrane domains or short C-terminal domains that interact with membrane-associated proteins such as Rab GTPases (Rab2-, 6-, 19-, and 30), Arf-like 1 (Arl1), or GRASP family proteins^13,16–18^. Initial vesicle capture results from interactions between the variable N-terminal domain and proteins on the vesicle surface or in the case of GMAP-210 direct interaction with vesicle lipids via its amphipathic lipid packing sensor (ALPS) motifs^19, 20^. Following long range capture, the vesicles are brought close to the target membrane for fusion. Rab GTPase–binding sites within the coiled-coil region likely contribute to this process, perhaps by inducing folding of the coiled-coil at unstructured breaks in the alpha-helices or by the transfer of vesicles from the N-terminal to the Rab-binding domain on the same or an adjacent golgin^16,21–23^. Vesicles are then handed off to multi-subunit tethering complexes, such as the COG complex, and SNARE proteins that drive membrane fusion^24,25^.

In *Toxoplasma*, we have an incomplete understanding of the proteins that regulate Golgi structure and trafficking. A conserved COG complex was recently characterized, that functions with the p115/Uso1-like tethering factor to mediate intra-Golgi membrane trafficking^4,26 ,27^. Several components of the TRAPP III tethering complex have also been localized at the Golgi^28^, and other tethering complexes COVERT and HOPS function at the endosome-like compartments^29^. To date, golgin-like proteins with long coiled-coil domains have yet to be identified in *T. gondii*.

In this study, we characterize a previously unstudied golgin-like protein, TgME49_213392, in *Toxoplasma gondii*. This protein localizes to the Golgi and contains a predicted ∼1,000-amino-acid coiled-coil domain, reminiscent of mammalian golgins. We demonstrate that this protein is essential for maintaining Golgi cisternal stacking and therefore named it TGIF (Toxoplasma Golgi Integrity Factor). Loss of TGIF results in severe defects in parasite replication and ultimately leads to parasite death. Furthermore, using a fluorescence-based pulse–chase assay, we show that disruption of Golgi structure in TGIF-deficient parasites impairs the trafficking of microneme and rhoptry neck proteins, whereas trafficking of rhoptry bulb proteins remains largely unaffected. These findings suggest that rhoptry bulb proteins may traffic through a Golgi-independent pathway. Collectively, this work advances our understanding of Golgi biology and the mechanisms regulating secretory protein trafficking in *T. gondii*.

## Results

### Identification of putative Rab6-associated proteins using yeast-2-hybrid

To identify new Golgi associated proteins, a yeast-2-hybrid screen was carried out to find putative interactors of Rab6, a GTPase located at the TGN^30^. Seven proteins were identified in the screen that had a confidence score of very high (A), high (B), or good (C) (**Table S1**). Since our interest was in Golgi-associated proteins, we focused on hits predicted to be essential for parasite survival, localized at or around the Golgi, or had a predicted structure or domain organization that suggested a role in trafficking. Using these criteria, two uncharacterized proteins were prioritized for further investigation: TgME49_313910 and TgME49_213392. A third protein, beta-COP (TGME49_266990), also fit these criteria but given the established role of this protein as a component of the vesicle coatomer COPI in other species, it was not examined in this study^31^.

We investigated the localization of TgME49_313910 by endogenously tagging the protein with a Ty-epitope. This protein did not co-localize with markers for the cis-Golgi (GRASP-mcherry)^32^ or the ER (GFP-HDEL)^33^. Instead, this protein had a punctate distribution throughout the cytosol (**Fig. S1A**). Due to this localization, TgME49_313910 was not investigated further, and we focused our efforts on characterizing TgME49_213392.

TgME49_213392 is a 254kDa protein that, based on InterPro analysis^34^, is predicted to contain an N-terminal RGPR-related domain (panther PTHR13402), typically found in the Sec16 family of proteins (amino acids (a.a.) 8-222). In addition, the protein is predicted to contain an intrinsically disordered region (IDR) spanning amino acids 135-865 (**Fig. 1A; purple)**. Analysis using coiled-coil prediction software N-coils and Marcoils revealed a long coiled-coil domain, although the predicted start and end of this region varied (a.a.1104-2153 and a.a.1146-2079 respectively) (**Fig. 1A grey, and S1B**). The predicted Rab6-binding site is located within this region (a.a 1124-1554) (**Fig. 1A**). After the coiled-coil, there is a short tail domain (**Fig. 1A, dark blue**). Given that Sec16 family proteins are essential for organizing ER exit sites and facilitating COPII vesicle formation in mammalian cells, and that the long coiled-coil domain is reminiscent of Golgin-family proteins, we further investigated the structural features of this protein.

**Figure 1:**
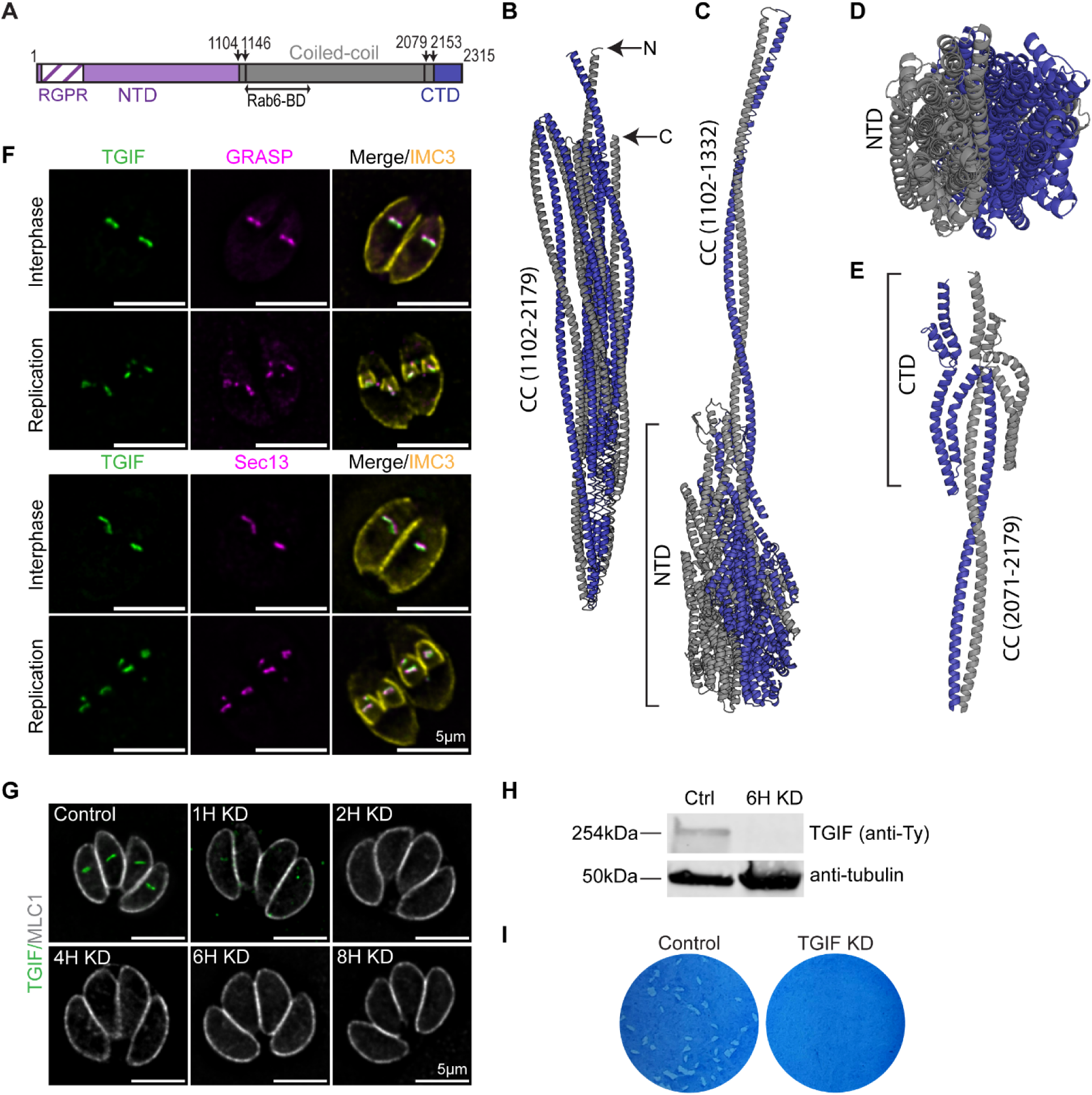
TGIF is an essential protein localized to the cis-Golgi. (A) Schematic illustration of TGIF domains predicted using Interpro, Marcoil and N-coils. The N-terminal domain (NTD: M1-L1101) contains a predicted RGPR-like domain (a.a. 8-222) with the remaining predicted to be intrinsically disordered. N-coils and Marcoils predict different start and end positions of the coiled-coil domain (grey) (indicated with black arrows). The C-terminal domain is shown in blue. The Rab6 binding region is predicted to be between (a.a. 1124-1554). (B) Alphafold predicted model of dimeric TGIF coiled-coil domain (a.a. 1102-2179). Six long stretches of alpha-helices are broken by 5 short unstructured breaks leading to a folded confirmation. N and C-termini are indicated with arrows (C) Alphafold predicted model of the TGIF N-terminal domain (a.a. 1-1101) and the first coiled-coil segment (a.a. 1102-1332). The NTD is made up of tighly packed alpha helices. This is a low confidence prediction with pIDDT scores between 50 and 70. (D) Construct shown in (C) rotated 90°. (E) Alphafold predicted model of TGIF tail domain as a dimer with the last coiled-coil segment. The CTD is formed from four short alpha helices which together form a H shape. (B-E) One monomer is shown in dark blue and the other shown in grey. (F) Immunofluorescence of TGIF-Ty parasites showing colocalization between TGIF (anti-Ty; green) and the cis-Golgi markers mGRASP - mCherry (magenta, top) and Sec13-EmGFP (magenta, bottom) during interphase and replication. Parasite periphery is indicated using anti-IMC3 antibodies (yellow). Scale bars are 5μm. Representative images from three experimental replicates (N = 47 vacuoles). Images are single z-slices. (G) Immunofluorescence of TGIF-Ty parasites after IAA treatment for 1, 2, 4, 6, and 8 hours. Anti-Ty (green) was used to visualize TGIF and an anti-MLC1 antibody (grey) indicate parasite periphery. Scale bars are 5μm. Representative images from two experimental replicates (N > 20 vacuoles per condition). Images are maximum intensity projection of z-stacks. (H) Western blot with anti-Ty (TGIF) and anti-tubulin (loading control) on TGIF-Ty parasite lysates harvested after 6 hours of IAA treatment, along with control parasites treated with ethanol. (I) Plaque assay of TGIF-Ty parasites grown for 8 days with ethanol (control) or IAA (knockdown). No plaques form upon TGIF knockdown.

### Structural predictions of TgME49_213392 using AlphaFold

Given the presence of the predicted coiled-coil domain, we modeled TgME49_213392 as a dimer using AlphaFold3^35^ (**Fig. 1B-E**). Despite the prediction that the N-terminal domain was intrinsically disordered, this region is modeled as a globular structure made up of approximately 36 short, tightly packed alpha-helical segments with extensive interactions at the dimer interface (**Fig 1C and 1D**). However, this structure is predicted with low confidence (pIDDT <50). The tail domain (F2173-N2314) also has a series of short alpha-helices that form a H-shape in the dimeric molecule (**Fig. 1E**).

Residues 1104-2200 form an extensive coiled-coil region containing five breaks in the alpha helices (**Fig 1B).** Each coiled-coil segment has an average length of ∼255 (Å) (25nm), thus this molecule could extend to ∼150nm in length in a fully open confirmation. The breaks in the coiled-coil result in a predicted folded confirmation such that the N and C-terminal ends of the protein are in close proximity. In the cell, this molecule likely exists in dynamic equilibrium where an extended state could facilitate long range interaction with vesicles and folding to bring the vesicle in closer proximity with the cisternal membranes. Collectively, the domain organization of TgME49_213392 is indictive of the Golgin family of protein and was named TGIF for <u>T</u>oxoplasma <u>G</u>olgi <u>I</u>ntegrity <u>F</u>actor.

### TGIF is an essential protein localized at the Golgi

To determine TGIF’s localization and assess its function in the parasite, the protein was endogenously tagged with a 3xTy epitope and an auxin inducible degradation domain (mAID) at its C-terminus (**Fig. S1C**). Correct integration of the tag in this parasite line, hereafter referred to as TGIF-Ty, was confirmed by genomic PCR (**Fig. S1D**). To determine if TGIF localized to the Golgi, we modified the TGIF-Ty line by endogenously tagging Sec13, a marker of cis-Golgi with EmeraldFP (referred to as TGIF-Ty:Sec13-GFP) or by transiently expressed the cis-Golgi marker GRASP-mcherry (**Fig. S1E and S1F**)^36^. TGIF strongly co-localized with GRASP and Sec13 during both interphase and replication (**Fig. 1F**). Consistent with this, TGIF localizes adjacent to but not overlapping with SORTLR and GalNac-GFP, markers of the trans-Golgi network (TGN)^37^ (**Fig. S3A**).

To investigate TGIF’s function, we determined if IAA treatment effectively induced TGIF knockdown. TGIF-Ty parasites were treated with ethanol or IAA for varying lengths of time (1-8 hours), and immunofluorescence assays (IFA) and western blots were performed (**Fig. 1G and H**). In control parasites, TGIF has an elongated structure consistent with its Golgi association. After 1 hour of IAA treatment, TGIF is visible as a single punctum in 51% of parasites and undetectable in the remaining 49%. After 2, 4, 6, and 8 hours of IAA treatment, TGIF was undetectable by IFA (**Fig. 1G**). Western blot analysis showed complete loss of TGIF protein levels after 6 hours of IAA treatment (**Fig. 1H, Fig. S2A and S2B**). To confirm TGIF’s essentiality in the parasites, a plaque assay was performed. No parasite growth was detected after TGIF depletion, demonstrating the essentiality of this protein (**Fig. 1I**).

### TGIF is required for Golgi structure

Due to TGIF’s localization and structural similarity to mammalian Golgins, we investigated if TGIF played a role in maintaining Golgi structure. To evaluate Golgi morphology in the absence of TGIF, GRASP-mcherry was transiently expressed in the TGIF-Ty:Sec13-GFP parasite line. Intracellular parasites were treated with ethanol or IAA for 1, 2, 4, 6, 15, and 48 hours prior to fixation. In control parasites, the Golgi had the expected organization, an elongated structure apical to the nucleus (**Fig 2A and S3B**). After 1 hour of IAA treatment, the size of the Golgi, represented by GRASP and Sec13, was significantly reduced (61% and 47%, respectively), and appeared as a punctum rather than elongated structure (**Fig. 2A and B, S3B and C**). The punctum did not further decrease in size after longer IAA treatment times (2-15 hours) with IAA (**Fig. 2A and B; Fig. S3C**). When parasites were treated with IAA for 48 hours, GRASP appeared diffuse throughout the cytosol with no clear enrichment at the Golgi, and Sec13 was missing from approximately 20±5.9% of parasites (**Fig. 2A; Fig S3D**).

**Figure 2:**
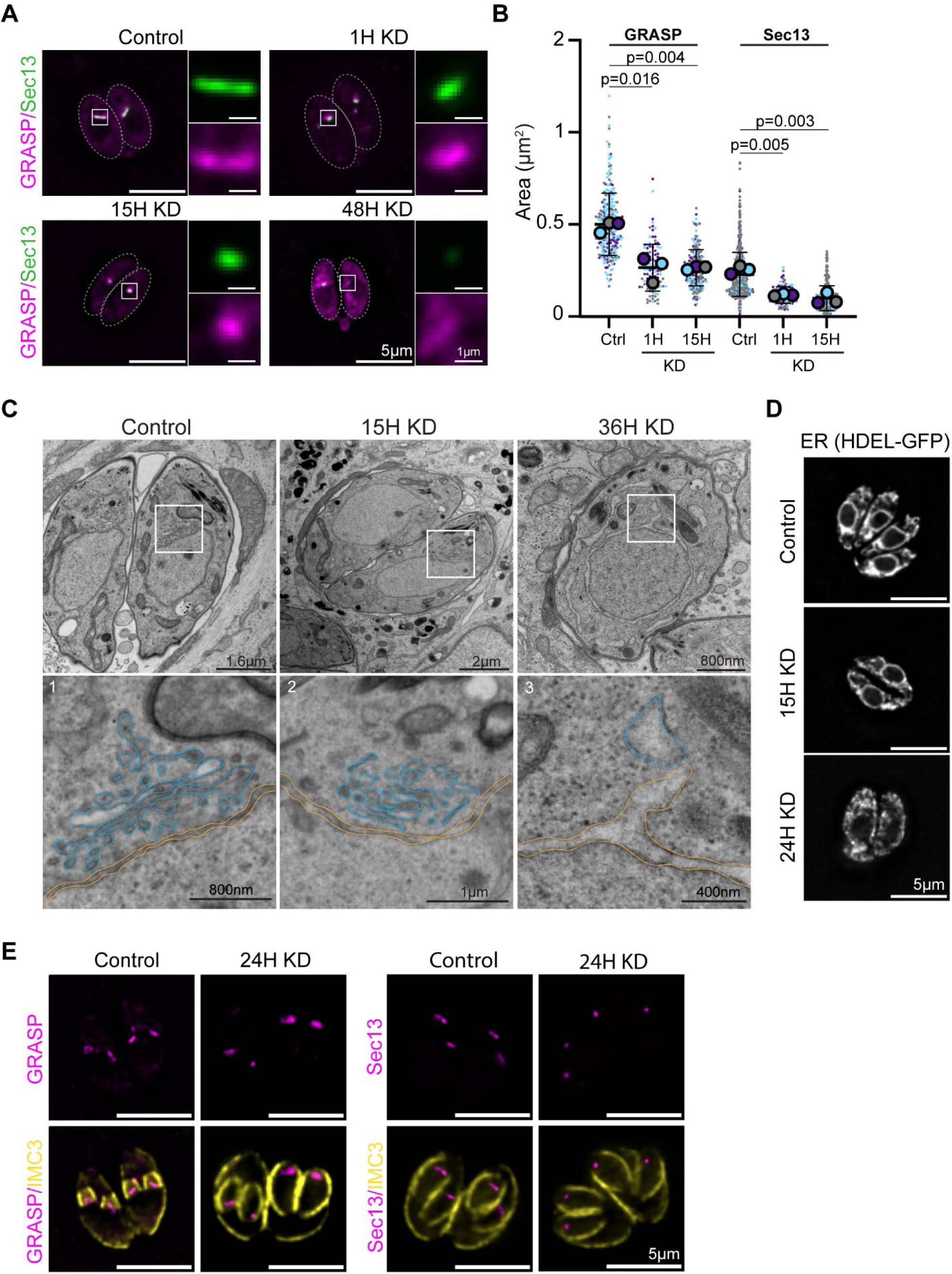
TGIF is needed for Golgi structural maintenance. (A) Immunofluorescence showing colocalization of mGRASP-mCherry (magenta) and Sec13-GFP (green) in TGIF-Ty:Sec13-GFP parasites treated with ethanol (control) or IAA (knockdown) for 1, 15, and 48 hours. White boxes denote region used to make insets. Scale bars are 5μm. Inset scale bar are 1μm. Representative images from three experimental replicates (N > 35 vacuoles per condition). Images are maximum intensity projections of z-stacked images. (B) Quantification of Golgi area from images in (A). P-values from paired Student’s t-tests are indicated. (C) Transmission electron microscopy images from TGIF-Ty parasites treated with ethanol (control) or IAA (knockdown) for 15 and 36 hours. White boxes denote region used to make insets. The Golgi cisternae are outlined in cyan, and the ER membrane is outlined in yellow in insets. Representative images from two experimental replicates (N > 12 vacuoles per condition). (D) Immunofluorescence showing the ER marker HDEL-GFP (grey) in TGIF-Ty parasites treated with ethanol (control) or IAA (knockdown) for 15 and 24 hours. Scale bars are 5μm. Representative images from two experimental replicates (N > 20 vacuoles per condition). (E) Immunofluorescence of mGRASP-mCherry (magenta, left) and Sec13-GFP (magenta, right) in TGIF-Ty:Sec13-GFP parasites treated with ethanol (control) or IAA (knockdown) for 24 hours. Anti-IMC3 (yellow) used to visualize the periphery of mother and daughter parasites. Scale bars are 5μm. Representative images from three experimental replicates (N > 30 vacuoles per condition).

To determine how the loss of TGIF disrupted Golgi structure at a higher resolution, we performed transmission electron microscopy (TEM) **(Fig. 2C**). In control parasites, elongated Golgi cisternae run parallel to the nucleus/ER (**Fig 2C, inset 1**). After 15 hours of IAA treatment, the cisternae were shorter, rounder and did not form an organized stack (**Fig. 2C, inset 2**). After 36 hours of IAA treatment, Golgi cisternae were no longer discernable, and the ER lumen appeared expanded (**Fig. 2C, inset 3**).

To further investigate the disruption to ER morphology, we transiently expressed GFP containing the ER retention sequence HDEL (HDEL-GFP)^33^ in TGIF-Ty parasites. In control parasites, the nuclear envelope is clearly visible with ER tubules throughout the cytosol (**Fig. 2D**). 15 hours of IAA treatment, the ER tubules in 13.4±3.4% of parasites were fragmented and nuclear envelope less defined. This percentage increased to 74±3.5% after 24 hours of IAA treatment (**Fig, 2D and S3E**). Collectively, this data indicates that TGIF is an essential protein necessary for cis-Golgi structural maintenance and loss of this protein has an indirect effect on ER morphology.

### Loss of TGIF does not disrupt Golgi division

Next, we sought to determine if loss of TGIF disrupted Golgi division or inheritance during cell division. Early in the *T. gondii* cell cycle, the Golgi associates with the duplicated centrosomes and splits twice by binary fission. Two Golgi fragments are then inherited by each daughter prior to fusion to form a single Golgi^38^. We treated parasites with ethanol or IAA for 24 hours and imaged vacuoles during division using IMC3 as a marker for daughter parasites. In control parasites, an elongated Golgi, denoted by GRASP and Sec13 markers, is inherited by each daughter parasite. Upon TGIF depletion, the Golgi were visible as single puncta, as seen in interphase, but TGIF depletion did not disrupt Golgi inheritance (**Fig. 2E)**. After 48 hours of IAA treatment, parasites frequently exhibited disrupted parasite morphology and asynchronous replication (**Fig. S3F).** Even with morphological defects, 90% of growing daughters contain Sec13 puncta indicating the loss of TGIF disrupted Golgi structure but not Golgi division or inheritance. Consistent with this, centrosome duplication was not disrupted by loss of TGIF (**Fig. S3G**). This data indicated that the disruption of Golgi morphology which results from loss of TGIF does not prevent Golgi segregation or inheritance into growing daughters.

### Loss of TGIF indirectly affects ER and TGN morphologies

Since TGIF was identified in a Rab6 Y2H screen, which is a marker of the TGN, we examined if loss of TGIF disrupted the trans-Golgi network organization, in addition to the cis-Golgi (**Fig. 2**). We examined the localization of three TGN markers: Rab6^39^, GalNac^40^, and SORTLR^41^ after 15 and 48 hours of IAA treatment. In control images, all markers are localized apical to the nucleus as expected, although GalNac localizes to a smaller punctum compared with SORTLR and Rab6, which were previously shown to colocalize^39^ (**Fig. 3A).** 15 hours after TGIF knockdown, no defect in TGN localization or organization was observed. However, 48 hours after TGIF depletion, we observed some knockdown parasites with normal morphology, while others had an altered morphology. At this time-point, the TGN marked by GalNac and SORTLR appears fragmented with structures localizing at the basal end of some parasites and absent in others, while Rab6 was diffuse throughout the cytosol (**Fig 3A**). These defects were observed in 69±5.9% and 43±9.2% of vacuoles, and 32±1.4% of parasites for GalNAc, SORTLR and Rab6, respectively. Given that disruptions in the cis-Golgi occur one hour after TGIF depletion and disruptions to the TGN markers are not observed until 48 hours after TGIF depletion, the TGN phenotypes are likely a downstream consequence of defects in cis-Golgi function, rather than TGIF having a direct role in TGN organization.

**Figure 3:**
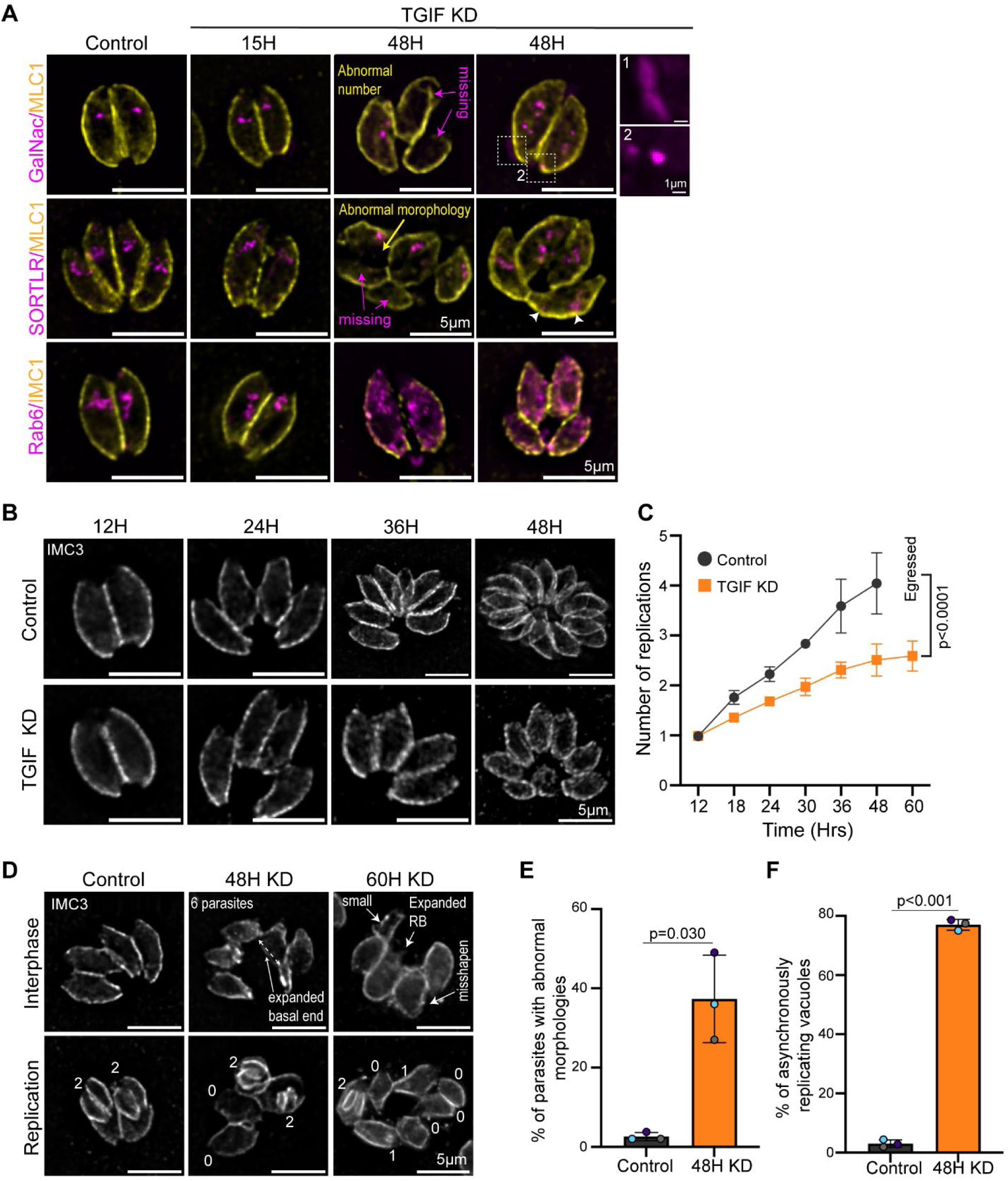
TGIF disrupts parasite replication. (A) Immunofluorescence showing the localization of trans-Golgi markers GalNac-GFP (magenta, top), SORTLR (visualized with an anti-SORTLR antibody, magenta, middle) and Rab6 (EmGFP-Rab6, magenta, bottom) in TGIF-Ty parasites treated with ethanol (control) or IAA (knockdown) for 15 and 48 hours. Anti-MLC1 or anti-IMC1 antibodies (yellow) used to visualize the periphery of parasites. 48 hours after TGIF knockdown, parasites exhibit both normal and abnormal morphologies (yellow arrow). Dashed white boxes denote area used to make insets to highlight mislocalization of GalNac in parasites with normal morphology. Magenta arrow indicates SORTLR and GalNac missing from parasites with abnormal morphology. Scale bars are 5μm and 1μm. Representative images from at least two experimental replicates (N > 20 vacuoles per condition). (B) Immunofluorescence showing parasite periphery labeled with IMC3 (anti-IMC3, grey) in TGIF-Ty parasites treated with ethanol (control) or IAA (knockdown) for 12, 24, 36, and 48 hours. Images are representative of the average number of parasites per vacuole at each time point. Scale bars are 5μm. Representative images from three experimental replicates (N > 100 vacuoles per time point per condition). (C) Graph shows parasite growth rate in control and TGIF knockdowns. After 60 hours of growth control parasites were extracellular. (D) Anti-IMC3 immunofluorescence showing abnormal parasite morphology in TGIF-Ty parasites treated with ethanol (control) or IAA (knockdown) for 48 and 60 hours in interphase (top) and replicating (bottom). In the bottom panels, the number of daughters per parasite are indicated. Scale bars are 5μm. Representative images from three experimental replicates (N > 100 vacuoles per time point per condition). (E-F) Quantification of the percent of parasites with abnormal morphologies, and percent of parasites replicating asynchronously after 48 hours of growth. Colored circles indicate averages from each independent replicates. P-values from paired Student’s T-tests are indicated.

Due to the defects in Golgi morphological observed in the absence of TGIF, we sought to determine if functions of the Golgi were disrupted, specifically glycosylation. The Golgi is home to glycosyltransferases which modify cargo proteins by O-glycosylation. TGIF-Ty parasites were grown with ethanol or IAA for 15 or 36 hours. Protein lysates were collected and western blots were stained with HPA conjugated to AlexaFluor-647 (*Helix pomatia agglutinin*) to recognize O-glycoproteins. Loss of TGIF disrupted O-glycosylation; proteins were observed with both increased (blue arrow) and decreased (red arrow) modification levels (**Fig. S4A**).

### TGIF affects parasite growth and replication

The Golgi contributes protein and lipids to the growing IMC, and having observed TGIF deficient parasites with altered morphology, we access this phenotype more closely. We first tested if TGIF is necessary for parasite replication. TGIF-Ty parasites were grown for 12-60 hours with ethanol or IAA and fixed for immunofluorescence at 6-hour intervals and then stained with anti-IMC3 antibodies, which marks the periphery of both mother and daughter cells. As expected, control vacuoles contained parasites that were a normal size, shape and formed the classic rosette morphology (**Fig. 3B),** while IAA treated cultures contained parasites with both normal and abnormal morphologies (**Fig. 3B and 3D**). To quantify if loss of TGIF affected the parasites’ replication rate, the number of parasites per vacuole was quantified for vacuoles with normal morphology. At the 18-hour time point, the growth rate of TGIF knockdown parasites was significantly slowed compared to control parasites. This trend continued at later time-points, with the TGIF growth plateauing with eight or less parasites per vacuole. By 60 hours, control parasites had egressed from the host cells, while TGIF KD parasites remained intracellular indicating a defect in natural egress (**Fig. 3C, S5A**).

Next, we examined vacuoles containing parasites with altered morphology. These phenotypes were first observed around 36 hours after IAA addition and increased in prominence at later time points. After 48 hours of IAA treatment, 37±6.4% of interphase parasites displayed morphological defects including expanded basal ends as well as smaller and misshapen cells. Parasites failed to form normal rosettes, and vacuoles contained an enlarged residual body (**Fig. 3D and 3E**).

In addition to these morphological abnormalities, synchronous replication of sister parasites was disrupted in 77±1.3% of replicating vacuoles after 48 hours of IAA treatment. In several instances, parasites within the same vacuole were at different stages of the cell cycle, or dividing parasites produced only a single daughter cell instead of two (**Fig. 3D and 3F**). Together, these data suggest that TGIF-mediated disruption of Golgi function impairs trafficking of proteins to the developing IMC during daughter cell formation.

### Induced egress and invasion were unaffected by the loss of TGIF

Given the inability of TGIF-depleted parasites to undergo natural egress, we next tested whether induced egress was also affected by the absence of TGIF. TGIF-Ty parasites were cultured for 36 hours in the presence of either ethanol or IAA, and differential interference contrast (DIC) microscopy was used to monitor egress following induction with the calcium ionophore, A23187. No significant difference was observed in the percentage of vacuoles undergoing egress between control and TGIF-depleted parasites. In both conditions, approximately 98% of parasites exited host cells within 5 minutes of ionophore treatment (**Fig. S5B, Videos S1 and S2**).

We also investigated if the loss of TGIF caused deficiencies in parasite invasion. TGIF-Ty parasites were grown with either ethanol or IAA for 24 hours. A 2-color invasion assay was performed^42^ and the percent of invaded parasites was calculated. Loss of TGIF did not disrupt parasite invasion (**Fig. S5C**).

### TGIF has an indirect effect on mitochondrial morphology

Cargo proteins that have dual localization in the apicoplast and mitochondria traffic through the Golgi, so we investigated if the localization or morphology of these organelles were disrupted in TGIF depleted parasites^43^. The mitochondrial marker Tim10 was transiently expressed in TGIF-Ty parasites and grown with ethanol or IAA for 15 and 48 hours. The mitochondria in control and TGIF depleted parasites (15 hours) had a lasso or linear “sperm-like” morphology as previously described^44^. In parasites grown for 48 hours with IAA, mitochondrial morphology was disrupted. The mitochondria was collapsed apical to the nucleus (**Fig. S5D**). To discern TGIF’s effect on the apicoplast, TGIF-Ty parasites were grown for 15 and 48 hours with ethanol or IAA and stained with anti-cpn60 antibodies. No apicoplast defects were observed when TGIF was depleted (**Fig. S5D**).

### Loss of TGIF disrupts processing and localization of microneme proteins

Microneme and rhoptry proteins are synthesized in the ER and are thought to traffic to the Golgi and then to post-Golgi compartments, which are comprised of a Rab5a-associated endosome-like compartment (ELC)^45^ and the PL-VAC^46^. We hypothesized that Golgi disruption due to the loss of TGIF would affect trafficking of secretory proteins. The localization of microneme proteins was investigated by expressing MIC8-mcherry in TGIF-Ty parasites or by immunofluorescence with antibodies that recognize AMA1, MIC2, proM2AP and M2AP (which recognizes both the mature (m) and unprocessed pro- (p) form of the protein) ^47–49^. In control parasites, proM2AP localized mostly to puncta at the apical end of the nucleus consistent with the expected localized at the ELC^50^ (**Fig. 4A, white arrow**). M2AP, MIC2, AMA1, and MIC8 all localized to the apical tip of control parasites, consistent with mature micronemes^50,51^ (**Fig. 4A and B, white arrows**). When TGIF was depleted for 24 hours, M2AP, MIC2, and MIC8 partially accumulated at the apical end of the nucleus, a localization consistent with the post-Golgi compartments (**Fig. 4A and B, arrowhead**), in addition to the mature micronemes at the apical tip (**Fig. 4A and 5B, arrow**). When TGIF was depleted for 24 hours, proM2AP puncta appeared more dispersed throughout the parasites, and staining was seen in the ER and residual body (RB) (**Fig. 4A)**. Interestingly, AMA1 localization did not change in TGIF depleted parasites (**Fig. 4B**, white arrow).

**Figure 4:**
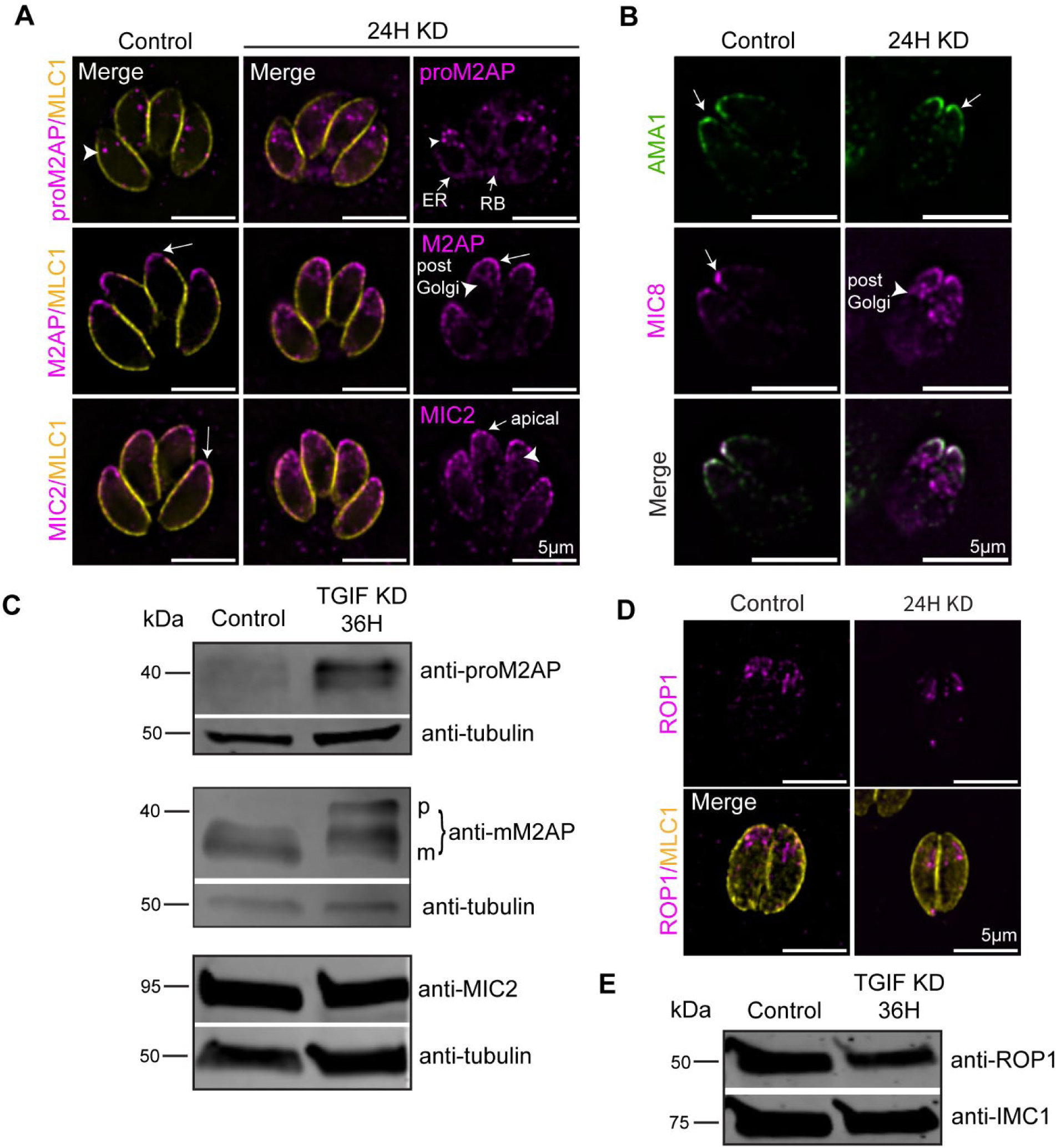
Microneme protein localization and processing is affected by the loss of TGIF. (A) Immunofluorescence showing the localization of microneme proteins proM2AP (anti-proM2AP, magenta, top row), M2AP (anti-M2AP, magenta, middle row), and MIC2 (anti-MIC2, magenta, bottom row) in TGIF-Ty parasites grown with ethanol (control) or IAA (knockdown) for 24 hours. Parasite periphery is indicated using anti-MLC1 antibodies (yellow). White arrowhead denotes the localization of proteins at the Golgi or post-Golgi compartments. White arrow shows protein localized at mature micronemes. Scale bars are 5μm. Representative images from two experimental replicates (N = 30 vacuoles per condition). (B) Immunofluorescence showing the microneme proteins AMA1 (anti-AMA1, green) and MIC8-mCherry (magenta) grown in TGIF-Ty parasites with ethanol (control) and IAA (knockdown) for 24 hours. White arrowhead denotes the localization of proteins at the Golgi or post-Golgi compartments. White arrow protein localization at mature micronemes. Scale bars are 5μm. Representative images from two experimental replicates (N = 20 vacuoles per condition). (C) Western blot with anti-proM2AP, anti=M2AP, anti-MIC2, and anti-tubulin (loading control) on TGIF-Ty parasite lysates harvested after 36 hours of ethanol or IAA treatment. Representative blots from three experimental replicates. (D) Immunofluorescence showing ROP1 fused to neonFP (magenta; top) grown in TGIF-Ty parasites with ethanol (control) or IAA (knockdown) for 24 hours. Parasite periphery is indicated using anti-MLC1 antibodies (yellow; bottom). Scale bars are 5μm. Representative images from three experimental replicates (N > 40 vacuoles per condition). (E) Western blot with anti-ROP1 and anti-IMC1 (loading control) on TGIF-Ty parasite lysates harvested after 36 hours of ethanol and IAA treatment. Representative blots from three experimental replicates.

**Figure 5:**
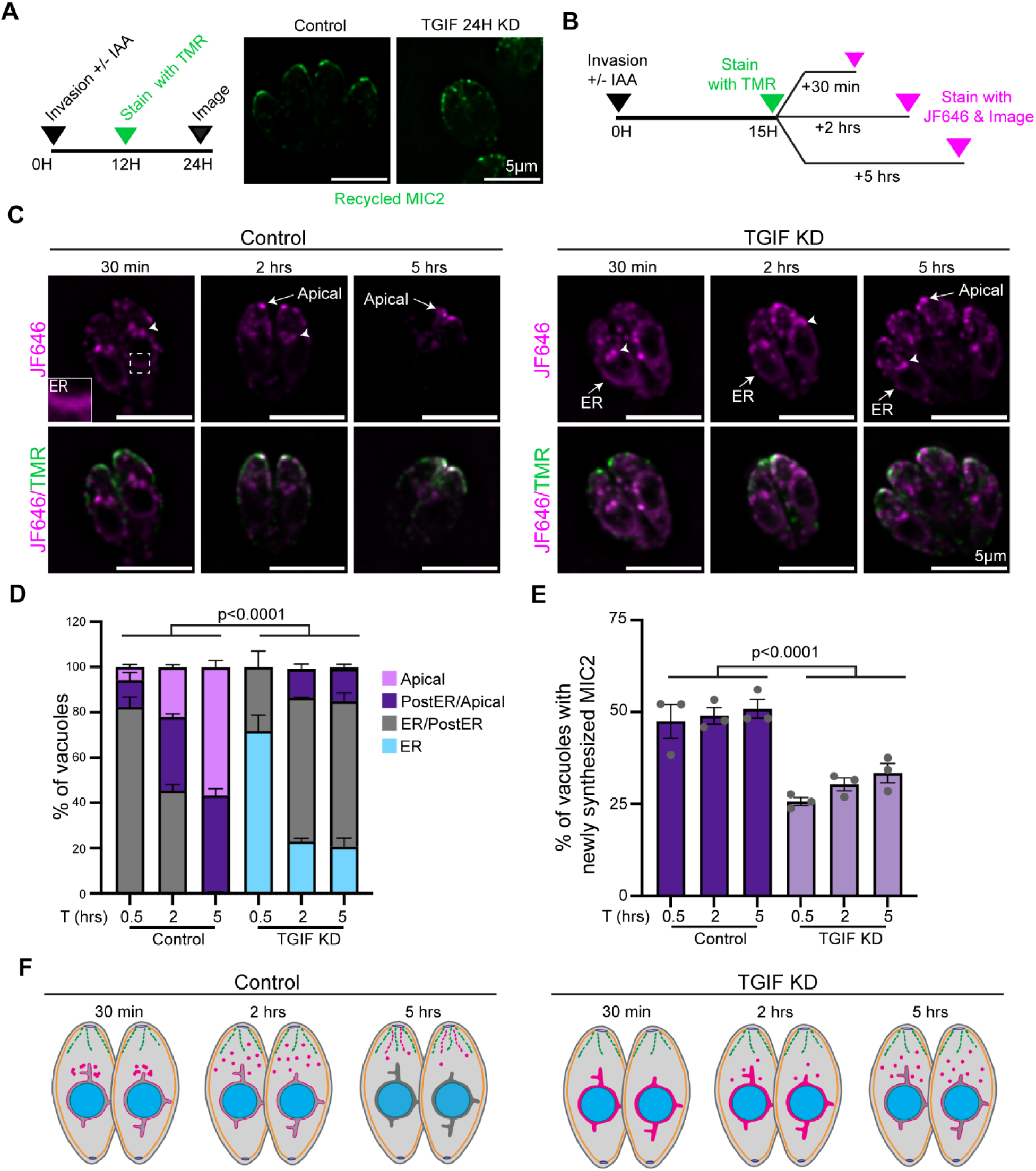
Loss of TGIF affects newly synthesized MIC2 trafficking. (A) *Left*. Schematic of the pulse-chase recycling assay. *Right.* Immunofluorescence showing recycled MIC2 stained with Halo-TMR (green) in TGIF-Ty:MIC2-Halo parasites grown for 24 hours with ethanol (control) or IAA (knockdown). Scale bars are 5μm. Representative images from three experimental replicates (N> 45 vacuoles per condition). (B) Schematic of the pulse-chase trafficking assay. (C) TGIF-Ty:MIC2-Halo parasites labeled with HaloLigand-TMR (green) and HaloLigand-JF646 (magenta) after growth with ethanol (control) or IAA (knockdown). Time points indicate period of growth between 1^st^ (TMR) and 2^nd^ (JF646) labeling reactions. Scale bars are 5μm. Representative images from three experimental replicates (N > 300 vacuoles per time point per condition). (D) Stacked bar chart indicating the localization of MIC2 labeled with JF646 at each time point. P-values are indicated. (E) Bar graph indicating the percent of vacuoles with JF646 staining at each time point, indicating the presence of newly synthesized protein. P-value indicated. (E) Illustration of MIC2 trafficking pathway in control and TGIF knockdown parasites.

To discern if TGIF depletion affected proteolytic processing or expression of microneme proteins, cell lysates from TGIF-Ty KD parasites were collected after growth in ethanol or IAA for 36 hours. Western blots were performed with antibodies against proM2AP, M2AP, and MIC2 (**Fig. 4C**). The levels of proM2AP increased by 45±2.2% when TGIF was depleted compared to control parasites (**Fig. 4C and S2C and F)** and the levels of mature (m) M2AP decreased by 73±0.06% when TGIF was depleted (**Fig. 4C and S2D and F**). MIC2 protein levels did not change between control and TGIF depleted parasites (**Fig. 4C and S2E and F**).

Since MIC8 trafficking is dependent on Rab5a which localizes to the ELC, we investigated if loss of TGIF affects the localization of Rab5a^51^. Transient expression of Rab5a in TGIF-Ty parasites grown for 15 and 48 hours in either ethanol or IAA revealed that loss of TGIF does not affect Rab5a localization at the ELC (**Fig. S6A**).

Next, we investigated if the localization of rhoptry proteins was affected by the loss of TGIF. TGIF-Ty parasites transiently expressing ROP1-neon were grown for 24 hours with ethanol or IAA. ROP1 localization at the apical end of parasites was not affected by the loss of TGIF (**Fig. 4D**). To discern if TGIF depletion affected the expression or processing of ROP1, we performed western blots using an anti-ROP1 antibody on TGIF-Ty lysates that were collected after being grown for 36 hours with ethanol or IAA (**Fig. 4E**). ROP1 protein levels decreased by 30±0.04% in TGIF-depleted parasites but ROP1 proteolytic processing was not affected (**Fig. S2G and S2H**). Together, this data indicates that loss of TGIF does not have a large impact on either rhoptry localization of expression.

### Development of a pulse-chase assay to track formation of secretory vesicles

Immunofluorescence assays indicated that loss of TGIF had a modest or minimal effect on the micronemes and rhoptries, respectively. However, these organelles are formed through both *de novo* synthesis and inheritance during parasite cell division ^52,53,54^. To specifically address if one or both of these pathways were disrupted upon TGIF depletion, we adapted a microscopy-based pulse-chase assay^52^. For this, three new parasites lines were created where the microneme protein MIC2, rhoptry bulb protein ROP1 and rhoptry neck protein RON2 were endogenously tagged with Halo in the TGIF-Ty parasite^55^.

We optimized the halo dye labeling reactions using TGIF-Ty:MIC2-Halo parasite line. Parasites were grown overnight and stained with HaloTag-Ligand-TMR (Halo-TMR), this labeling reaction should saturate all Halo-tagged proteins in the cell so the staining with a second dye, JaneliaFluor646 HaloTag-Ligand (Halo-JF646), would stain only newly synthesized proteins. To check that this was the case TGIF-Ty:MIC2 were stained with Halo-TMR, washed, and then immediately stained with Halo-JF646 HaloTag Ligand and imaged. No parasites were stained with JF646, verifying saturation of the Halo protein with the 1^st^ (TMR) dye (**Fig. S6B and S6C**).

### Loss of TGIF affects newly synthesized MIC2 trafficking

To determine if loss of TGIF perturbed microneme recycling, TGIF-Ty:MIC2-Halo parasites were grown for 12-hours post-invasion, stained with TMR, and grown for a further 12 hours before imaging. TMR-labeled micronemes were observed in both control and knockdown parasites indicating that loss of TGIF did not disrupt microneme recycling (**Fig. 5A**). To determine if loss of TGIF affects trafficking of newly synthesized MIC2, TGIF-Ty:MIC2-Halo parasites were grown for 12 hours with ethanol or IAA, pulse-labelled with TMR, and grown for 30 minutes, 2 hours, or 5 hours before labeling with Halo-JF646 and immediately live imaging (**Fig. 5B**). At the 30-minute time point, 48±8% of parasites contained newly synthesized protein and in those, the majority (82%) contained newly synthesized MIC2 proteins in the ER (**Fig. 5C; inset**) and apical to the nucleus in a localization consistent with the Golgi or post-Golgi compartment (**Fig. 5C; arrowhead**). In the remaining parasites (18%), MIC2 had trafficked out the ER and was solely localized in the post-Golgi compartments or at the apical tips consistent with mature micronemes (**Fig. 5C-E**). At the 2- and 5-hour time points the percent of parasites containing newly synthesized protein was the same as at the 30-minute time point (49±4% and 51±5%, respectively). However, at these later time-points, newly synthesized MIC2 was localized at the post-Golgi compartments and at the apical tip 58% and 100% of vacuoles, respectively (**Fig. 5C-E**), indicating that MIC2 exited the ER and trafficked to apically localized micronemes.

When TGIF was depleted, there was a significant reduction in the percent of parasites containing newly synthesized protein (26±2%, 30±3% and 33±5% at the 30 minute, 2- and 5-hour time points, respectively). In addition, the trafficking of MIC2 protein was perturbed. At the 30-minute time point, 75% of vacuoles had MIC2 staining in the ER, with no staining solely in the Golgi, post-Golgi or apical regions of the cell (**Fig. 5C and D**). At the 2- and 5-hour time-points, MIC2 remained in the ER and ER/PGC regions in ∼80% of parasites (**Fig. 5C and D**). Collectively, this data demonstrates that the loss of TGIF results in a decrease in MIC2 protein synthesis and disrupts the trafficking of newly synthesized MIC2 to mature micronemes (**Fig. 5F)**.

### Loss of TGIF affects RON2 synthesis and trafficking

Next, we investigated the effects of TGIF depletion on RON2 recycling and trafficking as described above. 12 hours after TMR-labeling, RON2-TMR protein was visible in all cells indicating that the protein was inherited during cell division **(Fig. 6A**). When the trafficking assay was performed (**Fig. 6B**), 25±3% of control parasites contained newly synthesized RON2 protein at the 30-minute time point, and 76% of those had RON2 staining in the ER, with the remaining 24% having RON2 positive puncta that were localized at the apical end of the cell. Some of these puncta were in proximity to the nucleus (**Fig. 6C**; **arrowhead**) while others were at the apical tip of the parasite (**Fig. 6C and D, arrow**). Additionally, when RON2 was localized in the ER, parasites contained elongated or lobed nuclei indicative of parasites that were undergoing endodyogeny. The percent of apically localized RON2 increased to 55% and then 85% at 2 and 5 time points, respectively (**Fig. 6B and 6C**). The percent of parasites with newly synthesized RON2 increased to 37% at the 5-hour time point.

**Figure 6:**
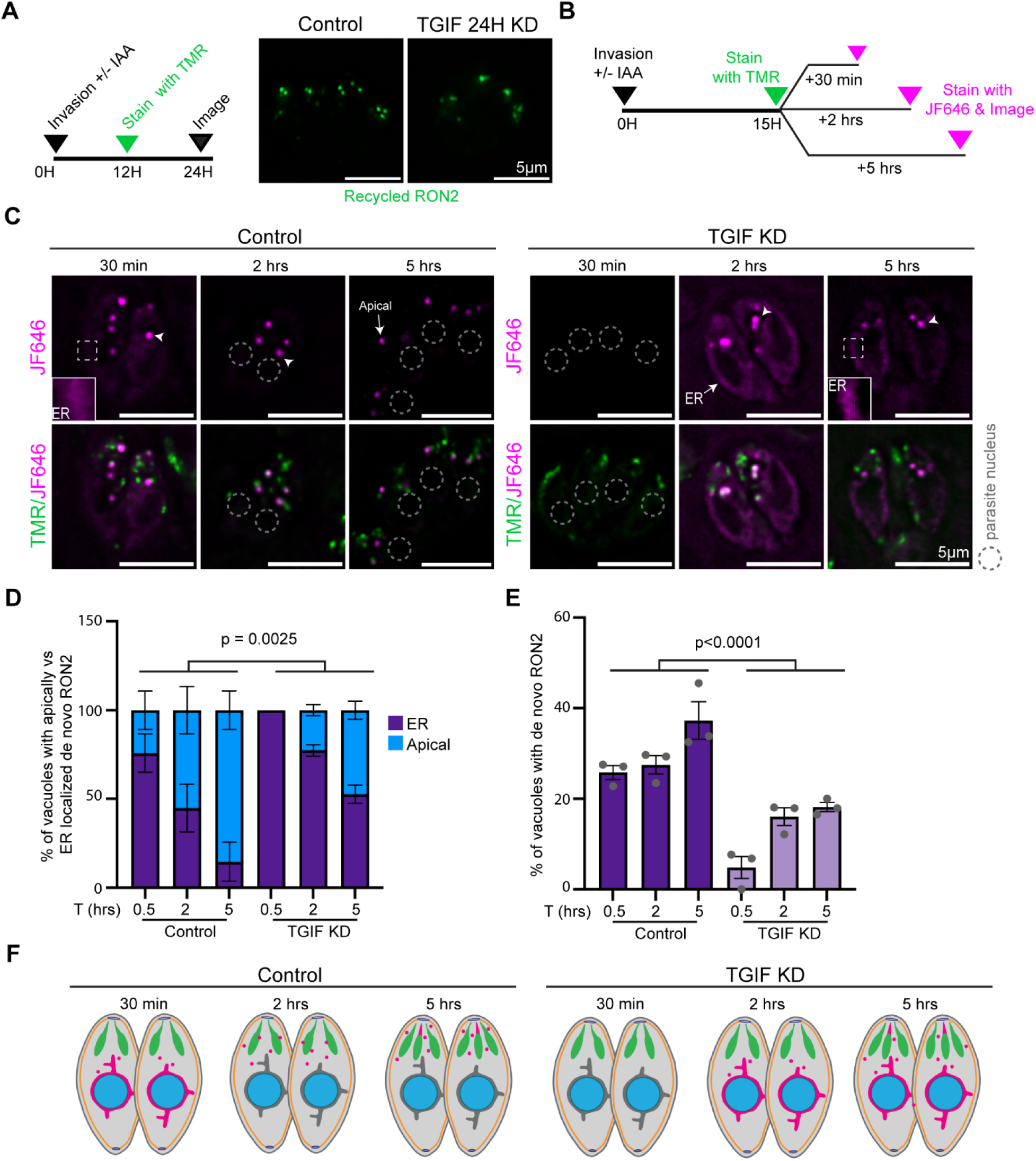
Loss of TGIF affects RON2 synthesis and trafficking. (A) *Left*. Schematic of the pulse-chase recycling assay. *Right.* Immunofluorescence showing recycled RON2 stained with Halo-TMR (green) in TGIF-Ty:RON2-Halo parasites grown for 24 hours with ethanol (control) or IAA (knockdown). Scale bars are 5μm. Representative images from two experimental replicates (N = 16 vacuoles per condition). (B) Schematic of the pulse-chase trafficking assay. (C) TGIF-Ty:RON2-Halo parasites labeled with HaloLigand-TMR (green) and HaloLigand-JF646 (magenta) after growth in ethanol (control) or IAA (knockdown). Time points indicate period of growth between 1^st^ and 2^nd^ labeling reactions. Scale bars are 5μm. Representative images from three experimental replicates (N > 300 vacuoles per time point per condition). (D) Stacked bar chart indicating the localization of newly synthesized RON2 at each time point from (C). P-value indicated. (E) Bar graph indicating the percent of vacuoles with JF646 staining at each time point, indicating the presence of newly synthesized protein. P-value indicated. (G) Illustration of RON2 trafficking pathway in control and TGIF knockdown parasites.

In contrast, when TGIF was depleted only 5±4% of vacuoles contained newly synthesized RON2 (**Fig. 6C and E**), when newly synthesized RON2 was observed, the protein was localized to the ER in all parasites. At the later time points, the percentage of parasites with newly synthesized RON2 increased to only 18±2% at the 5-hour time point. In contrast to what was observed in control parasites where 15% of parasites had ER localized RON2 at the 5-hour time point, 53% of parasites contained ER-localized RON2 after TGIF knockdown. Collectively, this data indicates that RON2 is expressed in a cell cycle dependent manner and disruption of TGIF affects both the trafficking and expression of the protein (**Fig. 6F**).

### Loss of TGIF disrupted rhoptry bulb maturation but not post-ER trafficking

When the recycling was performed with the ROP1-Halo line, we again observed that ROP1 recycling occurring during parasite cell division in both control and TGIF deficient parasites **(Fig. 7A**). When the trafficking experiment was performed **(Fig. 7B**), 35±10% of control parasites contained newly synthesized protein stained with Halo-JF646 (**Fig. 7C-7E)**. In contrast to what was observed with MIC2 and RON2, ROP1 was not observed in the ER at the 30-minute time point. Instead, ROP1 puncta were observed throughout the parasite’s apical end (**Fig. 7C, arrow; Fig. 7D**). At 2-hour hour time point, we observed puncta at the apical end that were often in alignment, and narrow tubules were observed emanating from the puncta (**Fig. 7C; 2 hours; inset**). In addition, canonical rhoptry bulbs, which appear as elongated structures of uniform thickness, were observed. The percentage of parasites containing canonical rhoptry bulb structures increased from 8% at the 2-hour time point to 50% at the 5-hour time point, while the percentage of parasites containing newly synthesized ROP1 increased about 80% at the 5-hour time point (**Fig. 7D**).

**Figure 7:**
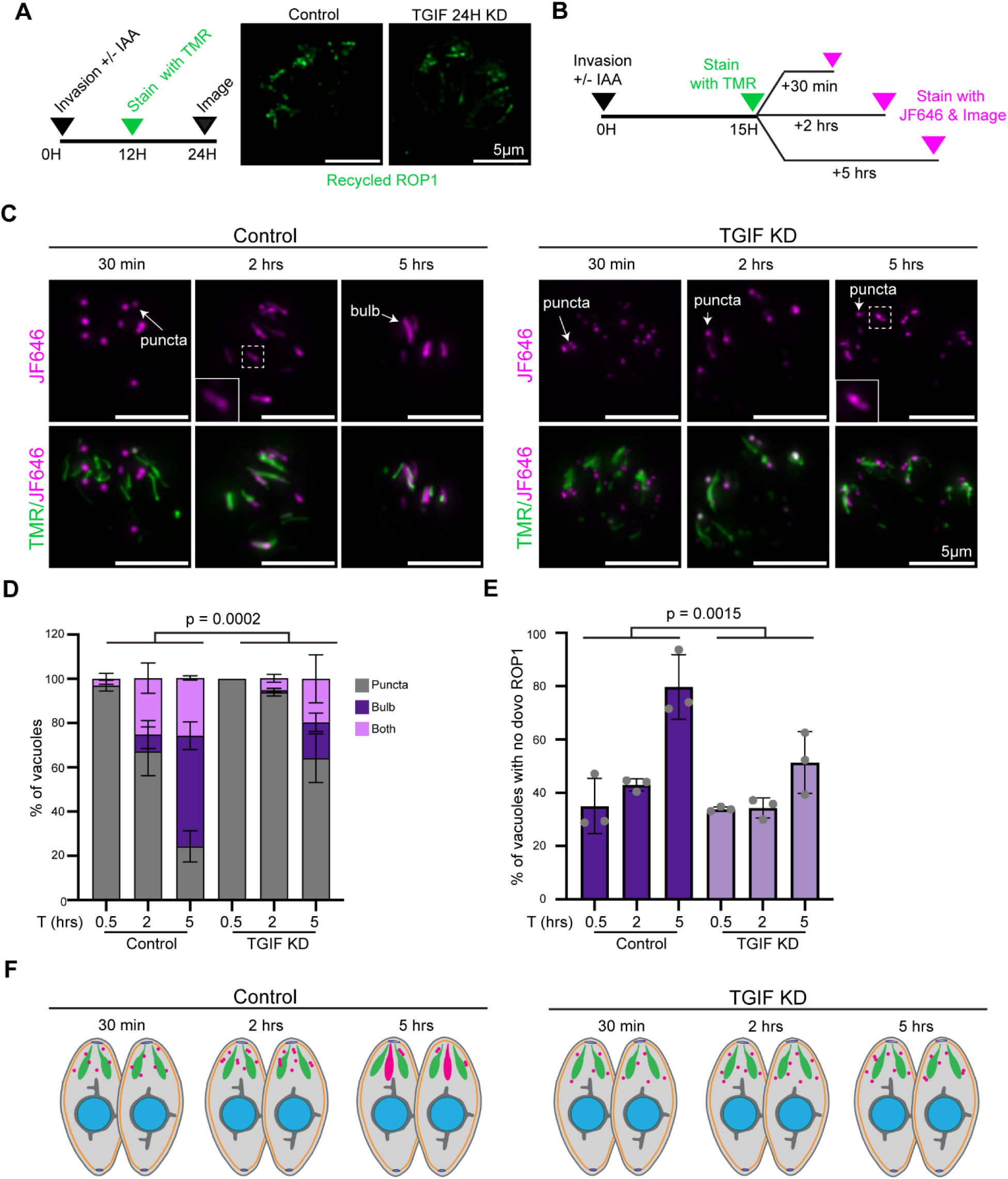
Loss of TGIF does not affect ROP1 synthesis or ER exit. (A) *Left*. Schematic of the pulse-chase recycling assay. *Right.* Immunofluorescence showing recycled ROP1 stained with Halo-TMR (green) in TGIF-Ty:ROP1-Halo parasites grown for 24 hours with ethanol (control) or IAA (knockdown). Scale bars are 5μm. Representative images from two experimental replicates (N = 17 vacuoles per condition). (B) Schematic of the pulse-chase trafficking assay. (C) TGIF-Ty:ROP1-Halo parasites labeled with HaloLigand-TMR (green) and HaloLigand-JF646 (magenta) after growth in ethanol (control) or IAA (knockdown). Time points indicate period of growth between 1^st^ and 2^nd^ labeling reactions. Scale bars are 5μm. Representative images from three experimental replicates (N > 300 vacuoles per time point per condition). (D) Stacked bar chart indicating the localization of newly synthesized ROP1 at each time point from (C). P-value indicated. (E) Bar graph indicating the percent of vacuoles with JF646 staining at each time point, indicating the presence of newly synthesized protein. P-value indicated. (G) Illustration of Rop1 trafficking pathway in control and TGIF knockdown parasites.

In TGIF depleted parasites, the percentage of parasites containing newly synthesized ROP1 was unchanged compared to control parasites (35±10% and 34±0.1 in control and knockdown parasites respectively) (**Fig. 7E**) and the localization of ROP1 was unchanged between knockdown and control, i.e. – puncta throughout the parasites apical end. At the 2-hour time point, differences in the morphology of ROP1 positive structures between control and knockdowns were observed. In the absence of TGIF, no parasites exclusively contained ROP1 bulbs and only 5% of parasites contain a mixture of bulbs and puncta in the absence of TGIF compared with 25% in controls (**Fig. 7D**). In addition, at the 5-hour time point, the percentage of parasites with newly synthesized ROP1 decreased to about 51% (compared with 80% in controls) (**Fig. 7E**)

Taken together these data show that 30 minutes after synthesis ROP1 is found in fluorescent puncta throughout the apical end of the parasites and was not observed in the ER, it’s well-established site of synthesis which indicates rapid ER exit^56–58^. Between 2 and 5 hours after synthesis, ROP1 is found in mature rhoptry bulbs at the parasite’s apical end. Surprisingly, loss of TGIF does not appear to disrupt ROP1 ER exit but disrupts the maturation of rhoptry bulbs (**Fig. 7F**).

## Discussion

The Golgi apparatus plays a central role in the trafficking and post-translational modification of proteins within the endomembrane system. Although *Toxoplasma gondii* possesses a simplified Golgi architecture compared to metazoans—consisting of a single Golgi stack—our understanding of the proteins that govern its organization and trafficking functions remains incomplete. In this study, we identified and characterized a Golgi-associated protein that is essential for maintaining Golgi structure and function, which we term TGIF (Toxoplasma Golgi Integrity Factor). TGIF is composed of an extended ∼1000-amino-acid coiled-coil domain, a globular N-terminal domain, and a short C-terminal tail. This domain organization is similar to Golgin family proteins known to mediate long-range vesicle capture and tethering at the Golgi.

### TGIF is a Golgin-like protein

AlphaFold structural predictions of TGIF indicate that its extensive coiled-coil domain is organized into six long α-helical segments, each approximately 250 nm in length, separated by five short unstructured regions. When fully extended, this coiled-coil region could theoretically span up to ∼1.5 µm from the cisternal surface. However, the structural model suggests that these helices adopt a folded, accordion-like conformation. In vivo, TGIF likely exists in a dynamic equilibrium between extended and compact states, potentially modulated by interactions with vesicles or other binding partners.

In contrast to many golgin-family proteins, which typically possess short N-terminal domains^59^, TGIF contains a large N-terminal region comprising roughly half of the protein. Although this domain is predicted to be intrinsically disordered by InterPro, AlphaFold modeling of a dimeric construct suggests it may adopt an elongated, globular structure composed of tightly packed short α-helices; however, this prediction is associated with low confidence. The C-terminal tail consists of four short α-helices arranged in an “H”-like configuration, with the final coiled-coil segment connecting with the crossbar. We hypothesize that this compact C-terminal region mediates association with Golgi membrane associated proteins.

### TGIF is an essential protein required for Golgi structure and trafficking

Using the auxin-inducible degradation system, TGIF protein levels were depleted within two hours of IAA treatment, resulting in rapid perturbations in Golgi morphology. Fluorescence microscopy revealed that the cis-Golgi, visualized using GRASP and Sec13, transitioned from an elongated structure at the apical end of the nucleus to a single punctum. Transmission electron microscopy further demonstrated shortening and disorganization of Golgi cisternae (**Fig. 2**). Following the initial disruption observed after approximately one hour of IAA treatment, no additional changes in Golgi morphology were detected over the subsequent fifteen hours, and Golgi inheritance remained intact during this period. At later time points (∼36–48 hours after IAA addition), the Golgi was absent in ∼20% of parasites, indicating a progressive loss of Golgi cisternae.

Despite the disorganization of cis-Golgi structure, depletion of TGIF did not affect the positioning of the trans-Golgi network (TGN) or the downstream endosome-like compartment marked by Rab5 within the first ∼36 hours following protein loss. However, after 48 hours, the strict apical localization of TGN markers was disrupted, with TGN fragments observed at the basal end in ∼50% of parasites (**Fig. 3**). At this stage, Rab6 also exhibited a diffuse cytosolic localization. The delayed onset of TGN defects relative to cis-Golgi disruption suggests that these later phenotypes are likely secondary consequences of impaired cis-Golgi function, rather than reflecting a direct role for TGIF in maintaining TGN organization.

TGIF represents the first Golgin-like protein identified in *T. gondii*, and its essentiality distinguishes it from Golgins in other eukaryotes, where multiple family members localize to distinct Golgi cisternae. In those systems, loss of individual Golgins typically results in mild phenotypes, whereas combinatorial knockouts produce progressively severe defects, consistent with functional redundancy^60^. The essential nature of TGIF may therefore reflect a limited repertoire of Golgin-like proteins in *T. gondii* or a reduced capacity for functional compensation.

Loss of Golgi cisternal organization in TGIF-depleted parasites coincides with defects in the trafficking of proteins destined for the inner membrane complex (IMC), micronemes, and the rhoptry neck. Given the tight coupling between Golgi structure and function, these phenotypes alone do not resolve the precise mechanistic role of TGIF, which will require further investigation. Based on its Golgin-like domain organization and its identification in a yeast two-hybrid screen as a potential Rab6 interactor, we hypothesize that TGIF may function as a vesicle tether at the Golgi. Future studies will focus on comprehensive characterization of the TGIF interactome, determining whether TGIF and Rab6 interact directly, and defining the functional consequences of this interaction. Notably, the Golgi defects observed following TGIF depletion differ from those reported upon disruption of the conserved oligomeric Golgi (COG) complex, a multi-subunit tethering complex^4^. Depletion of COG3 or COG7 subunits results in Golgi vesiculation, whereas loss of TGIF leads to a reduction in Golgi size and cisternal disorganization. These phenotypic differences do not exclude a role for TGIF in vesicle tethering, as they may reflect differences in cargo specificity or trafficking directionality (anterograde versus retrograde). Alternatively, TGIF may serve a structural role in maintaining Golgi morphology and cisternal stacking. Mechanisms governing Golgi organization in *T. gondii* remain poorly understood. Although the parasite encodes a single GRASP homolog, this protein is not thought to be essential for parasite survival^38^. Identifying proteins that interact with the N- and C-terminal domains of TGIF will be critical for elucidating its mechanistic role in Golgi organization and function.

### IMC trafficking pathways and natural egress were disrupted upon loss of TGIF

Upon exit from the Golgi, cargo proteins traffic to multiple subcellular destinations although the specific trafficking pathways are incompletely understood. Our data indicates that loss of TGIF differentially affects cargo trafficking. The most prominent phenotype observed following TGIF knockdown is a disruption in inner membrane complex (IMC) formation, accompanied by a reduced rate of parasite replication. These defects suggest that TGIF is required for efficient trafficking of IMC-associated proteins through the Golgi. Similar replication phenotypes, characterized by abnormal parasite morphology and arrest prior to the eight-parasite-per-vacuole stage, have been reported following disruption of Rab11b and clathrin heavy chain, both of which localize to the trans-Golgi network (TGN) further supporting the conclusion that TGIF depletion impairs trafficking of proteins required for IMC biogenesis during replication. However, the defects observed upon TGIF loss are less severe than those resulting from disruption of Golgi-associated syntaxin complex components, including SLY1, Stx6, and Vps45, which lead to more pronounced replication arrest^61^. These phenotypic differences may reflect distinct functional roles. TGIF may act as an initial vesicle tether within the Golgi, improving the efficiency of trafficking between cisternae, rather than directly mediating vesicle fusion events, as is characteristic of syntaxin-family proteins. Additionally, defects in IMC biogenesis that occur in the absence of TGIF may arise indirectly from perturbations in ER morphology, which likely occur as a downstream consequence of Golgi disruption as a recent study have demonstrated that the ER contributes lipids to IMC formation through membrane contact sites^62^.

Upon arrest of replication at the eight-parasite-per-vacuole stage, parasites failed to undergo natural egress. However, calcium ionophore-induced egress remained intact, suggesting that sufficient microneme secretion capacity was retained. This observation indicates that loss of TGIF likely affects signaling pathways upstream of intracellular calcium mobilization rather than the execution of egress itself.

Several secreted dense granule proteins are known to regulate parasite egress. These include diacylglycerol kinase 2 (DGK2), which converts DAG to phosphatidic acid—a key signaling molecule controlling natural egress^63^, lecithin:cholesterol acyltransferase (TgLCAT), which generates lysophosphatidylcholine from phosphatidylcholine^64^ and TgGRA41^65^ although the mechanisms by which TgLCAT and TgGRA41 influence egress remain unclear. While we were unable to apply the Halo-based pulse-chase approach to dense granule proteins due to difficulties in endogenously tagging GRAs with large epitopes, the observed defects in microneme and rhoptry trafficking raise the possibility that dense granule protein trafficking is also impaired in the absence of TGIF.

A second possibility is a disruption in calcium signaling, a key regulator of parasite egress. Loss of the ER localized calcium channel TgSERCA also results in delayed natural egress as SERCA activity is required to maintain intracellular Ca²⁺ stores^66^. Notably, loss of TGIF resulted in pronounced ER disorganization, fragmentation, and expansion of the ER lumen, as observed by transmission electron microscopy. These defects in ER architecture may further contribute to the impaired natural egress phenotype observed in TGIF-depleted parasites.

### Microneme and rhoptry proteins exhibit different trafficking pathways

To investigate how TGIF loss affects microneme and rhoptry trafficking, we first examined the localization of these organelles, which arise from a combination of inheritance during cell division and de novo synthesis^53^. Direct immunofluorescence assays revealed only partial mis-localization of microneme proteins. Therefore, to specifically track newly synthesized proteins, we employed a Halo-based pulse-chase assay to assess the trafficking of MIC2, RON2, and ROP1. This approach revealed key differences in their trafficking pathways.

Thirty minutes after assay initiation, newly synthesized MIC2 and RON2 was predominately localized to the ER, with additional staining near the Golgi or post-Golgi compartments. At later time points, the proportion of parasites with ER-localized protein decreased, while the proportion with apically localized protein increased. This pattern indicates that MIC2 and RON2 exit the ER, accumulate at Golgi or post-Golgi compartments (consisting of ELC or PL-VAC) and are delivered to their respective organelles (**Fig. 8; blue and yellow pathways**).

**Figure 8:**
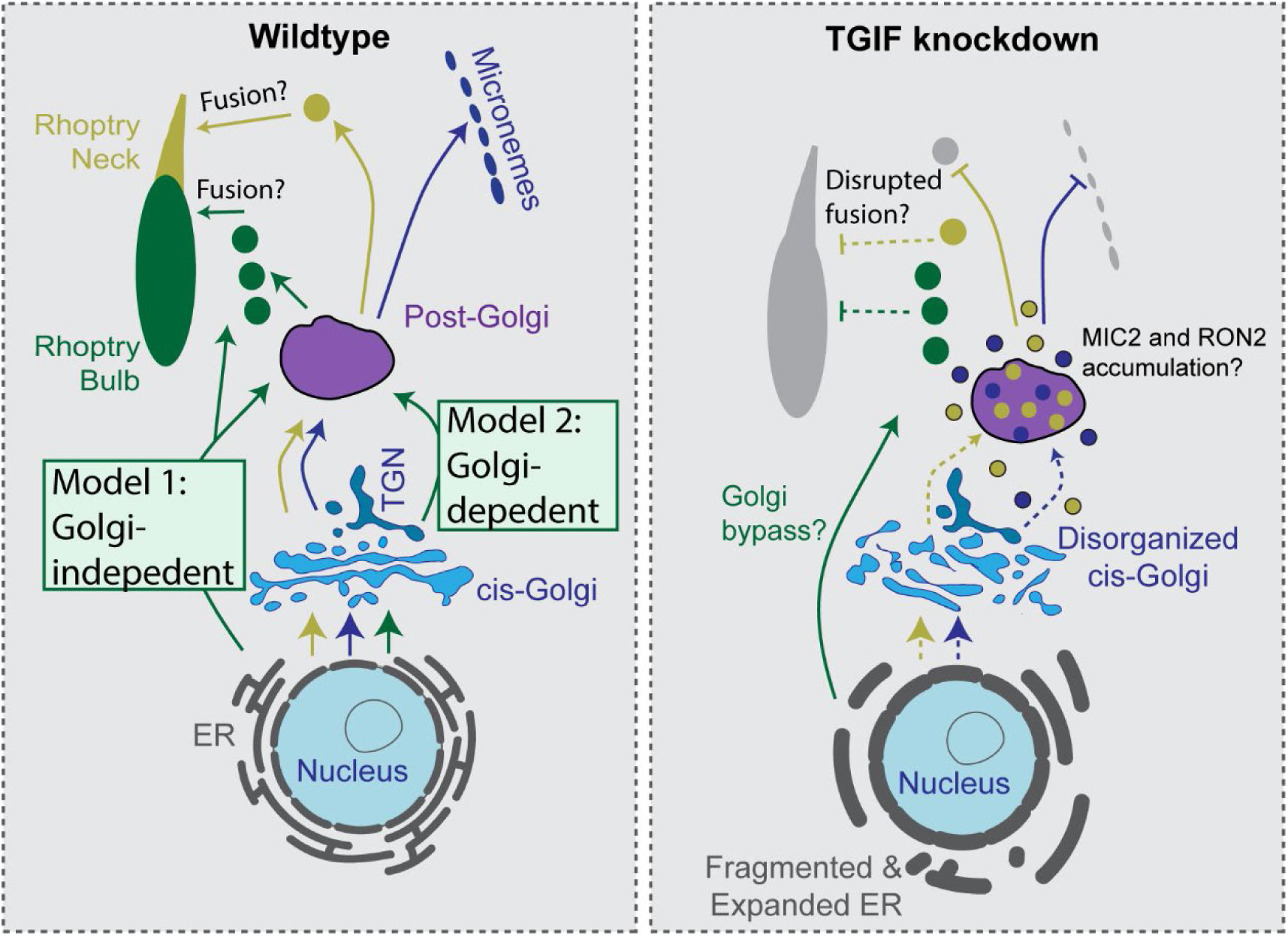
Model of TGIF function in microneme and rhoptry trafficking. Working model of the secretory pathway in *T. gondii*. After ER synthesis, MIC2 and RON2 traffic through the Golgi to the Post-Golgi compartments which comprise the PLVAC and Rab5a ELC (purple) and transport vesicles. Proteins are then found in micronemes (dark blue) and in the rhoptry neck (yellow). After synthesis ROP1 (green) rapidly exists the ER. We propose two potential models for ROP1 trafficking. In model 1, ROP1 uses a Golgi bypass pathway in wildtype parasites. ROP1 vesicles are trafficked from the ER either directly to the apical end or through the post-Golgi compartment(s). We propose that mature rhoptry bulbs form from fusion vesicles containing rhoptry bulb and rhoptry neck proteins. Alternatively, in model 2, ROP1 traffics through the cis- and trans- Golgi and then to the post-Golgi compartment before trafficking in vesicles to the apical end for maturation in wildtype parasites. When TGIF is depleted, ROP1 exit from the ER is not disrupted indicating it traffics via a Golgi-independent pathway. Once at the apical end, bulb formation is perturbed, suggesting the trafficking of factors necessary for bulb maturation is disrupted. Loss of TGIF also results in decreased synthesis of MIC2 and RON2 and accumulation of newly synthesized proteins in post-Golgi compartments or adjacent vesicles. Expanded ER lumen and disorganized cis-cisternae are observed in the absence of TGIF. Arrows indicate trafficking pathways; ROP1 in green, MIC2 in blue, RON2 in yellow. Dashed arrow indicates decreased trafficking. Blunt arrow indicates blocked trafficking pathways.

We observed a significant reduction in the percent of parasites containing newly synthesized protein upon loss of TGIF (35% and 51% reduction for MIC2 and RON2, respectively at the 5-hour time point). In both cases, newly synthesized protein was retained in the endoplasmic reticulum (ER) for several hours in contrast to control cells. Collectively, indicating that both the synthesis and trafficking of MIC2 and RON2 are impaired when TGIF is depleted and Golgi morphology is disrupted.

In contrast, ROP1 displayed a distinct trafficking and expression profile. Newly synthesized ROP1 was never observed in the ER at the 30-minute time point its established site of synthesis^56–58^. Moreover, ER exit of ROP1 was not disrupted, as more than 95% of parasites exhibited ROP1 puncta in both control and TGIF-depleted conditions at the 30-minute time point. These results could be interpreted in two ways. First, ROP1 could traffic in a Golgi-independent manner contrary to previously held assumptions. The rapid ER exit is consistent with a Golgi bypass pathway, which is typically faster than classic trafficking pathways^67^ (**Fig. 8; model 1**). Alternatively, ROP1 may traffic through the Golgi in wildtype cells but in the absence of TGIF a Golgi bypass pathway may be activated (**Fig. 8; model 2**). The prevailing understanding that rhoptry proteins trafficking in a Golgi dependent manner was based on studies using knockdown or dominant-negative parasite lines in which disruption of various proteins impaired rhoptry trafficking, formation, or proteolytic processing, as assessed by immunofluorescence or western blotting. Notably, these proteins include several factors localized to the trans-Golgi network (TGN) and endosome-like compartments, such as SNARE proteins, Rab1B, clathrin heavy chain and its associated adaptors, components of the HOPS/CORVET tethering complexes, and vacuolar proton ATPases at the PL- VAC ^61,46,68–71^. These studies are not necessarily inconsistent with the Golgi bypass model (model 1) as it may be the case that upon ER exit ROP1 traffics directly to the ELC, alternatively components necessary for rhoptry organelle maturation may trafficking in a Golgi-dependent manner. Golgi bypass pathways operate in both stress-induced and steady-state conditions in other cell types^67^ and distinguishing between these models will be the focus of future studies.

### Insights into rhoptry biogenesis

The two subdomains of the rhoptry (neck and bulb) have distinct proteomic composition that is sequentially secreted from the parasite during invasion^72^. The process of how this organelle forms and maintains protein organization in these subdomains is poorly understood. The pulse-chase assay reveals new insights into this process. In control parasites ROP1 positive puncta align at the parasite’s apical end and narrow tubule extensions were sometimes visible **(Fig. 7).** Bulb-shaped structures typical of mature rhoptries were then observed. Although speculative, we hypothesize that fusion of multiple ROP1 vesicular puncta with RON2 puncta leads to the formation of mature organelle. Although loss of TGIF did not disrupt ROP1 exit from the ER, rhoptry biogenesis was disrupted in that ROP1 puncta did not mature into bulb-shaped structures. The reason for this disruption is unclear. One possibility is that the maturation process depends on the accurate trafficking of rhoptry neck proteins, which are trafficked in Golgi-dependent pathway and disrupted upon TGIF knockdown. Future studies will focus on further characterizing these trafficking pathways and the molecular mechanisms governing rhoptry biogenesis.

In summary, TGIF is the first Golgin-family protein identified in *T. gondii*. It has essential roles in Golgi structural maintenance and trafficking and a pulse-chase assay in the TGIF knockdown lines provides new insights into the Golgi’s role in trafficking secretory cargo proteins.

## Materials and Methods

### Cell culture, drug selection, and inducible knock down

#### Cell culture and parasite transfection

*T. gondii* tachyzoites derived from the RH strain were used in all experiments. Parasites were maintained in human foreskin fibroblasts (HFFs) in Dulbecco’s Modified Eagle Media (DMEM) (ThermoFisher, Carlsbad CA) containing 1% (v/v) heat inactivated fetal bovine serum (FBS) (VWR, Radnor PA), 1X antibiotic/antimycotic (ThermoFisher), 1mM sodium pyruvate (Gibco Cat#11360-070) as described in Roos et al., 1994^73^ and hereafter referred to as DMEM growth media. Cells were grown at 37°C, 5% CO_2_. All parasite transfections were completed as described in ^74^ using a BTX electroporator with the following settings: voltage 1500V, resistance 25Ω and capacitance 25μF. For transient transfection, 25μg of plasmid was transfected and grown in confluent HFF monolayers on either MatTek dishes (MatTek corporation, Ashland MA) for live cell imaging or on coverslips before fixed cell imaging. For CRISPR-Cas9 tagging, 25ug of pU6 plasmid containing Cas9 and appropriate guideRNA was transfection along with 50ul homologous recombination (HR) oligo produced by PCR with Q5 polymerase as per manufactures instructions (NEB).

#### Parasite drug selection and inducible knock down

To make TGIF-Ty-mAID parasite line, transfect parasites were selected for the presence of HXGPRT gene using DMEM media containing 25μg/mL mycophenolic acid (MPA, Sigma Catalog#: M5255-50MG) and 50μg/mL xanthine (Sigma Catalog#: X0262-5G) which was added to cells 24 hours after transfection. To induce knock down using the AID system, parasites expressing TGIF-Ty-mAID were treated with 25μM of 3-indoleacetic acid (IAA/auxin) (Sigma-Aldrich Catalog#: I2886) ^75^.

### Yeast-2-Hybrid of Rab6

A yeast-2-hybrid screen was performed by Hybrigenics Services as previously described ^76^. Constitutively active TgRab6 (TgME49_310460) (Rab6 Q70L) lacking the last three amino acids (CSC) to prevent palmitoylation and membrane association was cloned into the pB27 vector (N-LexA-bait-C fusion) and transformed into yeast. The Rab6 construct was screened against the RH cDNA library for interactions. The confidence scores for each interaction were evaluated and determined algorithmically (Predicted Biological Score, PBS). Results were filtered to exclude any interactions with antisense or out-of-frame prey proteins.

### Parasite line creation

All primers, antibodies and plasmids used in this study are listed in Tables S2, S3 and S4 respectively.

#### Creation of TGIF-Ty parasite line

pTKOII_MyoL_3xTymAID (unpublished plasmid) was digested with AvrlI and BglII to remove the MyoL genomic DNA sequence. TGIF genomic DNA was obtained by PCR using genomic (g) DNA obtained from RH-Tir1 parasites ^75^ and primers TGIF gF1 and TGIF gR2. Plasmid backbone and insert were gel purified and ligated via Gibson assembly using NEBuilder HiFi DNA assembly master mix (New England BioLabs, Ipswich MA). Plasmids were transfected in NEB5α bacteria, clones were screened by PCR, and positive clones were verified by Sanger sequencing. 25μg of pTKOII_TGIF_3xTymAID plasmid was transfected into Tir1 parasites and parasites were drug selected using MPA and xanthine. Parasites were subcloned by limiting dilution to a 96-well plate (Corning Catalog#: 3585) containing a confluent monolayer of HFFs and grown for 8 days. Positive clones were confirmed by fluorescence microscopy staining with anti-Ty antibody and genomic integration confirmed by PCR using primers F1, R1 and R2. A single clone was chosen and used for all future experiments.

#### Creation of a TGIF-Ty: Sec13_EmGFP parasite line

To endogenously tag Sec13 in TGIF-Ty parasites using CRISPR-Cas9, TGIF-Ty parasites were transfected with 25μg of plasmid DNA containing a Cas9 expression cassette with a guide RNA for Sec13, and a homologous recombination oligomer (HR oligo) produced by PCR using primers Sec13-GFP HR F1 and R1 (45μL). The Sec13 guide RNA was designed using the gRNA design tool on Benchling (https://benchling.com). Sec13 proteospacers (primers) were reconstituted to 200µM in nuclease free water. Primers diluted to 100nM in 1X T4 ligation buffer (NEB Cat# B0202SVIAL). To anneal the proteospacers, samples were heated to 95°C for 5 minutes, and cooled slowly to 4°C at a rate of 0.1°C/second. A Cas9 containing plasmid was digested with Bsa1 and the duplexed Sec13 protospacers was ligated into the plasmid backbone using T4 DNA ligase (NEB Cat# M0202S), and incubated at RT for 2 hours. The plasmid was transformed into DH5⍺ competent *E. coli* cells (NEB Cat#C2987I). Mini-prepped using a QIAprep kit (Cat# 27106), were confirmed by sequencing before transfecting into parasites^77^. The primers for the HR oligo were designed to amplify GFP and had overhangs complementary to the last 40bp of sec13 (forward primer) and the first 40bp of the 3’ UTR (reverse primer). Parasites were grown for 2 days before FAC sorting for GFP positive parasites and subcloned into a 96-well plate. Positive clonal lines were confirmed by immunofluorescence and PCR verification using primers SG-F1, SG-R1 and SG-R2. A single clone was chosen and used for all future experiments.

#### Creation of TGIF-Ty:MIC2-Halo, TGIF-Ty:RON2-Halo, and TGIF-Ty:ROP1-Halo

Plasmids containing guideRNAs for MIC2, RON2, and ROP1 in a pU6_Cas9_YFP plasmid and a plasmid containing Halo coding sequence were generously gifted to us by Drs Markus Meissner and Simon Gras. To create each line, 25μg of plasmid DNA of either MIC2, RON2, or ROP1 guideRNAs and 45μL of Halo-HR oligo produced using PCR with Q5. were transfected into TGIF-Ty parasites. Parasites were grown for 2 days before being FACsorted for YFP positive parasites and subcloned into a 96-well plate. A single clone that had Halo-tagged protein expression, confirmed using HaloTag Ligand TMR staining, were chosen and used for all future experiments.

### Microscopy

Fluorescence and DIC imaging was completed on a DeltaVision Elite microscope system with an Olympus base with a 100 x 1.39 NA and 60 x 1.42 NA objective. This microscope has an environmental chamber heated to 37°C. This system is equipped with a scientific CMOS camera and DV Insight solid-state illumination module with the excitation wavelengths: DAPI = 390/18 nm, FITC = 475/28 nm, TRITC = 542/27 nm, and Alexa 647 = 632/22 nm). Single band pass emission filters had wavelengths: DAPI = 435/48 nm, FITC = 525/48 nm, TRITC = 597/45 nm, and Alexa 647 = 679/34 nm. Fluorescent and contrast imaging was also completed on a Nikon TI-2 microscope system with 100 x 1.45 NA, 60 x 1.42 NA, and 40 x 0.60 NA objectives. This microscope has an environmental chamber heated to 37°C. This system is equipped with an ORCA-

Fusion C14440 digital CMSO camera and Lumencor Spectra light engine with excitation wavelengths: DAPI = 390/22 nm, FITC = 475/28 nm, Cy3 = 555/28 nm, and Cy5 = 637/12 nm. Single band emission filters had wavelengths: DAPI 432/36 nm, FITC = 515/30 nm, Cy3 = 595/31 nm, and Cy5 = 680/24 nm. A Z-step of 0.02μm was used to obtain multiple focal planes for fixed imaging. Fluorescent images used in figures are represented as a maximum intensity projection of Z-stacks of deconvolved images Brightness and contrast for all fluorescent images were adjusted for print. All adjustments were normalized within each data set.

#### Live-cell imaging

Parasites were grown in DMEM growth media with either ethanol or IAA. DMEM was replaced with Imaging media (Fluorobrite DMEM (ThermoFisher, Cat# A19967) containing 1% FBS and 1x antibiotic/antimycotic and prewarmed to 37°C). Images were acquired using a DeltaVision Elite microscope. Image acquisition speed was determined on a case-by-case basis and are indicated in the figure legends.

#### Fixed-cell imaging and immunofluorescent assays

Parasites were fixed with 4% paraformaldehyde (PFA) (Electron microscopy sciences, Hatfield PA, Cat# 15714) in 1xPBS (ThermoFisher, Cat# 18912-014) for 15 minutes at room temperature (RT). Parasites were washed 3x in 1xPBS and permeabilized in 0.25% TX-100 (ThermoFisher Cat# 28314) in 1xPBS for 15 minutes at RT. Parasites were washed 3x in 1xPBS and blocked in 2% BSA (ThermoFisher Cat# BP9703-100) for 15 minutes at RT before being washed 3x in 1xPBS. All primary and secondary antibodies were diluted in 2% BSA made in 1xPBS as indicated in Table 4. Parasites were incubated with primary and secondary antibodies for 30 minutes before being washed 3x in 1xPBS. DAPI was diluted in 1xPBS, and parasites incubated for 10 minutes before being washed 3x in 1xPBS. Coverslips were mounted on slides using either Prolong Gold anti-fade (ThermoFisher, Catalog# P36930) or Prolong Diamond anti-fade reagent (ThermoFisher, Catalog# P36965). Coverslips were allowed to dry overnight, and clear nail polish was added to the edges of the coverslips and dried for at least 10 minutes before imaging.

#### Transmission electron microscopy (TEM)

Aclar was cut to fit into 2” shell vials. Aclar pieces were dipped into ethanol and dried completely before being placed into 6-wells and seeded with HFFs. Once HFFs were confluent, 3x10^6^ parasites were grown in DMEM containing ethanol or IAA for 15 or 36 hours. Parasites were washed with 1xPBS and fixed with 1.5mL EM fixative (2.5% glutaraldehyde in 0.1M Na cacodylate buffer, 3mM MgCL_2_, pH 7.4) for 1 hour at RT. Fixed parasites were delivered to the Bioscience Electron Microscopy Laboratory (BEML) facility at The University of Connecticut for EM preparation. Images were taken on a FEI Tecnai 12 G2 Spirit Bio TWIN transmission electron microscope equipped with an AMT NanoSprint12 (12-megapixel CMOS) camera.

### Assays

#### Plaque assay

HFFs were grown to confluence in 6-well plates (Corning, Glendale AZ, Catalog#: 3506). 200 TGIF-Ty parasites were added to each well and grown with ethanol or IAA for 8 days. For rescue plaque assays, parasites were grown with IAA for 2, 4, and 15 hours, the media containing IAA was washed out, replaced with normal DMEM media, and parasites were added to wells and grown for 8 days. Media was aspirated and replaced with 100% methanol pre-chilled to -20°C for 5 minutes. Methanol was removed and Coomassie stain (40% methanol, 7% acetic acid, 300μM Brilliant blue R [Sigma-Aldrich B7920]) was added for 2 hours followed by 1 hour of de-stain (40% methanol, 7% acetic acid). Images were taken on an iPhone.

#### Invasion assay

TGIF-Ty parasites split into two T25 flasks and grown for 24 hours in normal growth media, then treated with either ethanol or IAA for a further 24 hours. IAA treated parasites were scraped and syringe released as they did not naturally egress. 3x10^6^ parasites were added to coverslips containing a confluent HFF monolayer in DMEM growth media and allowed to invade for 1 hour. Parasites were fixed in 4% PFA + 0.8% glutaraldehyde in 1x PHEM (126mM PIPES, 25mM HEPES, 10mM EGTA, 8.2mM MgSO_4_, pH 7) for 15 minutes at 37°C. Glutaraldehyde was quenched in freshly made 1.25% sodium borohydride (w/v) in 1xPBS for 7 minutes. Coverslips were washed 3x in 1xPBS and stained with mouse anti-SAG1 (1:100) for 30 minutes at RT. Coverslips were washed 3x in 1xPBS and stained with goat anti-mouse AlexaFluor 488 (1:1000) secondary for 30 minutes. Coverslips were washed 3x in 1xPBS. Parasites were permeabilized with 0.25% TX-100 in 1xPBS for 15 minutes at RT and washed 3x in 1xPBS. Coverslips were blocked in 2% BSA for 15 minutes at RT and washed 3x in 1xPBS. Coverslips were stained with rabbit anti-IMC3 (1:1500, rb) for 30 minutes and wash 3x with 1xPBS. Coverslips were stained with goat anti-Rabbit AlexaFluor 546 (1:2000) secondary for 30 minutes and wash 3x with 1xPBS. Coverslips were mounted on slides with prolong gold and imaged using a 20x and 60x lens. 10 fields of view (2048X2048 pixel) randomly imaged. The number of intracellular and extracellular parasites was counted using the Cell Counter add-on in ImageJ and the percent of invaded parasites calculated. Graph in shown **Fig S5** is the result from three biological replicates. Total number of parasites counted in each experimental replicate was 829, 2955, and 3955.

#### Replication assay

2x10^6^ TGIF-Ty parasites were added to a 6-well with coverslips containing confluent HFFs. Parasites were grown in DMEM containing ethanol or IAA for 6, 12, 18, 24, 36, 48, and 60 hours. An IFA was completed as stated above. Parasites were incubated with anti-Ty (1:1000, ms) and rabbit anti-IMC3 (1:1500) made in 2% BSA. Parasites were incubated with goat anti-mouse 488 (1:1000) and goat anti-rabbit 546 (1:2000). 10 images were taken from randomly chosen fields of view using the 2048X2048-pixel field of view and 60x lens. The number of parasites per vacuole were counted (N > 300 vacuoles). To determine growth rate, no abnormal vacuoles were counted. Growth rate was calculated by taking the log_2_(average parasite number per vacuole). The combined results from three biological replicates are shown. Vacuoles containing parasites with abnormal morphologies were excluded from the growth rate analysis. The number of vacuoles containing parasites with abnormal morphologies were quantified separately.

#### Egress Assay

3x10^3^ TGIF-Ty parasites were added to MatTek dishes containing confluent HFFs and grown with DMEM containing ethanol or IAA for 36 hours. Normal growth media was replaced with 1.5mL of fluorobrite DMEM imaging media and temperature equilibrated at 37°C for 30 minutes before transferring to the microscope. DIC images were taken using the 60x objective lens. Images were taken for 10 seconds before adding 500μL of 20mM calcium ionophore (final concentration 5μM) (Sigma-Aldrich Cat# A23187) Images were taken every second for 5 minutes or until all vacuoles in the frame egressed. 7 separate control experiments and 10 separate TGIF KD experiments were completed with at least 2 Matteks per condition in each experiment. A total of over 204 control vacuoles and 273 TGIF KD vacuoles were counted per condition over these experiments.

### Halo Assays

#### Halo ligand control

TGIF-Ty:MIC2-Halo parasites were added to MatTek dishes containing confluent HFFs and grown with 1% DMEM media for 24 hours. 1% DMEM media was removed and cells incubated in 100µl of DMEM growth media containing 25µM containing HaloTag Ligand TMR (Halo-TMR) (Promega Cat# G8252) (1:200 dilution of 5mM stock) at 37°C for 15 minutes. Halo media was removed, parasites washed with 3x in 1% DMEM media. The labeling reaction was then repeated with 200µM JaneliaFluor646 HaloTag Ligand (Halo-JF646) (Promega Cat# GA1120) (1:1000 dilution of 5mg diluted to 200mM stock in H_2_O). Cells were washed with 3x in 1% DMEM and imaged live immediately in imaging media prewarmed to 37°C. One experiment was completed with 10 vacuoles imaged.

#### Halo recycling assay

TGIF-Ty:Halo-MIC2, TGIF-Ty:Halo-ROP1, or TGIF-Ty:Halo-RON2 parasites were added to MatTek dishes with ethanol or IAA and incubated for 12 hours. Labeling reaction with Halo-TMR was carried out as described above, washed with 3x in 1% DMEM media and grown for a further 12 hours in ethanol or IAA. DMEM growth media was replaced with 2mL of imaging media prewarmed to 37°C and imaged live. Two biological replicates experiments were completed for each parasite line with a minimum of 17 vacuoles per condition.

#### Halo trafficking assay

TGIF-Ty:Halo-MIC2, TGIF-Ty:Halo-ROP1, or TGIF-Ty:Halo-RON2 parasites were added to MatTek dishes containing confluent HFFs and grown with 1% DMEM media containing ethanol or IAA for 15 hours. Labeling reaction with Halo-TMR was carried out as described above, washed with 3x in 1% DMEM media and grown for a further 30 minutes, 2 hours, or 5 hours at 37°C in 1% DMEM media containing ethanol or IAA. At the end of each growth period, parasites were labeled with Halo-JF646 as described above. Cells were washed with 3x in 1% DMEM and imaged live immediately in imaging media prewarmed to 37°C. A total of three biological replicates were performed and over 300 vacuoles were imaged for each condition.

### Western blots

Parasite lysates were collected by spinning down freshly released or syringe released parasites from a T25 and centrifuged at 1,260 x *g* for 4 minutes. 6 x 10^7^ parasites were resuspended in 1x PBS. 10x sample buffer (0.6M Tris-HCl pH 6.8, 700 mM sodium dodecyl sulfate (SDS), 20% glycerol, 0.75 mM bromophenol blue, 2.5 mM 2-mercaptoethanol, and 25 mM dithiothreitol) was added to a final concentration of 2x and samples were boiled for 10 minutes at 95°C. Samples and protein ladder (Bio-Rad Catalog# 1610373) were run on 4-20% or 12% polyacrylamide gels (BioRad Catalog# 4561094, Catalog# 4561044). Proteins were transferred to a PVDF (polyvinylidene Fluoride) membrane (BioRad, Catalog# 162-0219) for 1 hour at 4°C. Membranes were blocked in blocking buffer (5% w/v milk powder in 1x TBST [0.1M Tris-base pH 7.4, 10% Tween-20, 0.15M NaCl, in water]) for 1 hours at RT with shaking. Primary and secondary antibodies were diluted in 5% w/v dry milk powder made in 1xTBS-T (**Table S4**). Primary antibodies were incubated overnight at 4°C with shaking, washed 3x with 1x T-BST and secondary antibodies were incubated at RT for 1 hour with shaking and washed 3x with 1x TBS-T. Blots were imaged on a Li-cor ODYSSEY CLx instrument. Briefly, for western blots probed for glycans, after protein transfer to the membrane, HPA-conjugated Alexa-647 (ThermoFisher Catalog#: L32454) used at 1:250 from a 1mg/mL working stock was diluted in Bio-Rab blocking buffer (Catalog# 12010020) and incubated over night at 4°C with shaking. Membranes were washed 3x in 1xTBS-T. Blots were imaged on a Li-cor ODYSSEY CLx instrument. Anti-a-tubulin was used as a loading control (1:5000) with a 1-hour RT incubation and probed with IR anti-mouse (LICOR bio-Catalog#: 926-68070) (1:1000) also incubated for 1 hour at RT before 3x washes with 1xTBS-T. Analysis was done using either ImageStudio or ImageJ2 v2.2.0/1.53t.

### Analysis

#### Image processing and analysis

Images were deconvolved using SoftWoRx v7.2.2 (Conservative ratio, 10 iterations). Images, including measurements and further processing were analyzed in ImageJ2 v2.2.0/1.53t. Statistical analysis was done using Microsoft Excel v16.69.1 or Prism GraphPad v10.6.1. Graphs were generated in Prism GraphPad v10.6.1.

#### Structural analysis using AlphaFold and Pymol

Protein sequences were obtained from ToxoDB (https://toxodb.org/toxo/app)^78^. ME49 strain sequences were used for structural analysis. Protein structures were obtained by submitting protein sequences into the AlphaFold 3 web-service. Structures from AlphaFold were uploaded into PyMOL (https://pymol.org/2/) for analysis. Coiled-coil prediction was done using the Marcoil and Ncoils programs on the Waggawagga prediction web-service (https://waggawagga.motorprotein.de/). Domain analysis was performed using Interpro (https://www.ebi.ac.uk/interpro/).

#### Statistical Analysis

Data analysis was performed in R using a likelihood ratio test to assess the effect of IAA-induced TGIF knockdown over time. Two linear models were made: a reduced model that included only time and replicates as factors, and a full model that included time, replicates, and treatment. These models were compared using an F test, and a p-value was derived from the F-statistic to determine if TGIF knockdown has a statistically significant effect on the overall time course. A three-way ANOVA with time, replicate, treatment and interaction between time and treatment was then performed. This method of data analysis was used to determine the P-value of the replication assay, and the time course assays with MIC2, RON2, and ROP1-halo. A p-value <0.05 was considered significant.

## Acknowledgements

We would like to thank members of the Heaslip Lab for helpful discussion during these experiments. We would like to thank Drs. Markus Meissner and Simon Gras (LMU, Munich, Germany) for sharing MIC2, ROP1, and RON2 CRISPR/Cas9 plasmids. Drs. Martiza Abril and Xuanhao Sun in the Bioscience Electron Microscopy Laboratory (University of Connecticut) for their help performing transmission electron microscopy and Dr. Amanda DelVichio at the UConn Flow Cytometry Core for her help FACS sorting parasite lines. We would like to thank Drs. Vern Cruthers (University of Michigan Medical School), Dr. Gary Ward (University of Vermont), Dr. Sabrina Marion (University of Lille), Dr. MJ Gubbels (Boston College), Dr. Chris de Graffenried (Brown University) and Dr. Boris Striepen (University of Pennsylvania) for generously sharing antibodies. We thank Drs. Adam Zweifach and Thomas Sladewski (University of Connecticut) for their assistance with statistical analysis and Pymol.

## Author Contributions Statement

CP and ATH contributed to the design of experiments. CP conducted the experiments. ATH and CP analyzed the data. Both authors co-wrote the paper.

