## Supplemental Data File for "TGIF is a Golgin-like protein required for Golgi structural maintenance and function in *Toxoplasma gondii*"

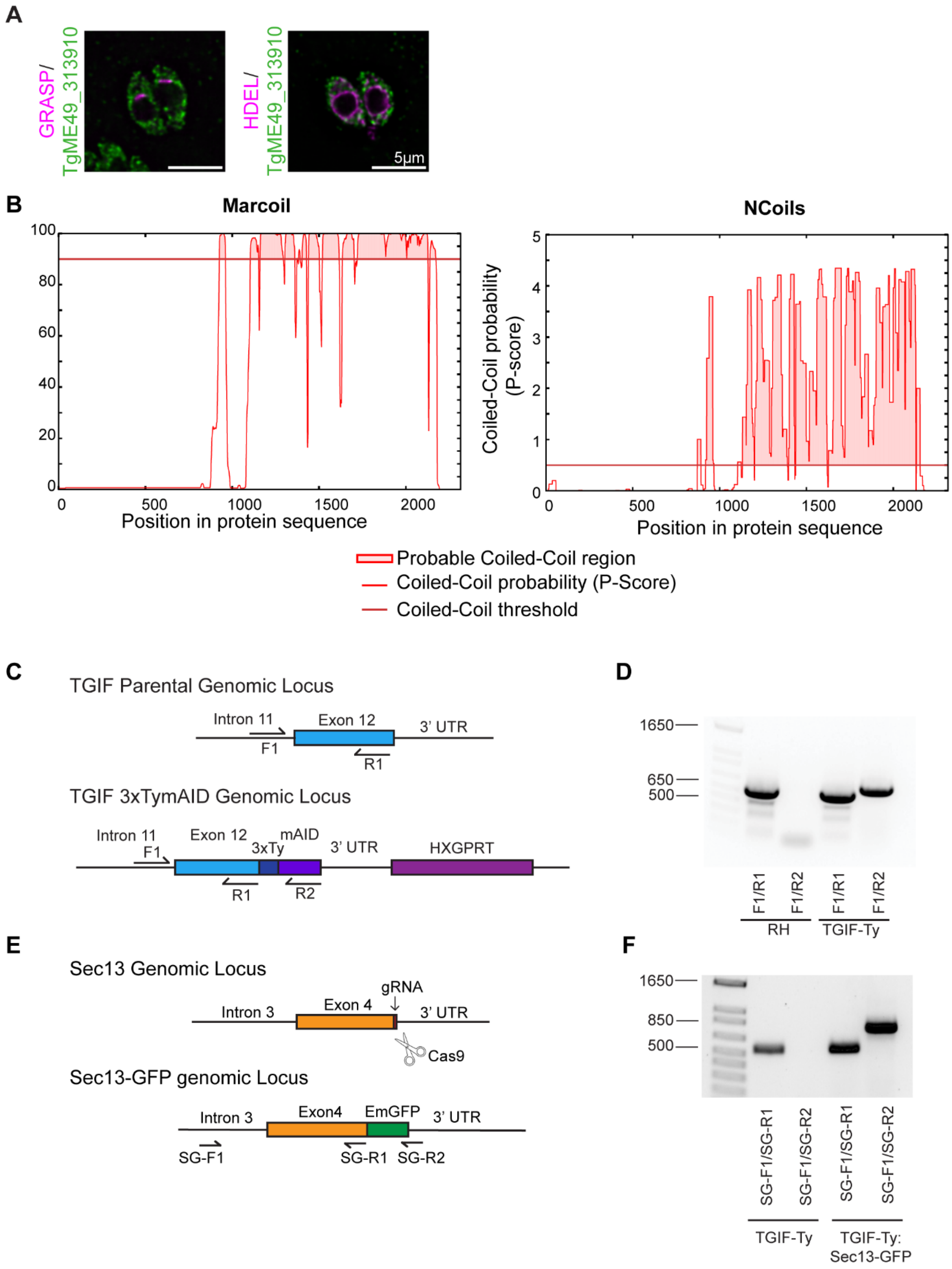
A
GRASP/
TgME49_313910
HDEL/
TgME49_313910
5μm
B
Marcoil
NCoils
Coiled-Coil probability
(P-score)
Position in protein sequence
Position in protein sequence
Probable Coiled-Coil region
Coiled-Coil probability (P-Score)
Coiled-Coil threshold
C
TGIF Parental Genomic Locus
Intron 11
Exon 12
3' UTR
F1
R1
TGIF 3xTymAID Genomic Locus
Intron 11
Exon 12
mAID
3xTy
3' UTR
HXGPRT
F1
R1
R2
D
1650
650
500
F1/R1
F1/R2
F1/R1
F1/R2
RH
TGIF-Ty
E
Sec13 Genomic Locus
gRNA
Intron 3
Exon 4
3' UTR
Cas9
Sec13-GFP genomic Locus
Intron 3
Exon4
EmGFP
3' UTR
SG-F1
SG-R1
SG-R2
F
1650
850
500
SG-F1/SG-R1
SG-F1/SG-R2
SG-F1/SG-R1
SG-F1/SG-R2
TGIF-Ty
TGIF-Ty:
Sec13-GFP

**Figure S1:** (A) Immunofluorescence showing the localization of TgME49_313910 using anti-Ty (green) and the cis-Golgi marker mGRASP fused to mCherry (magenta; left panel) or the ER marker (HDEL-GFP) (magenta; right panel). Scale bars are 5μm. Representative images from a single experimental replicate (N = 10 vacuoles for each marker). Images are deconvolved, single z-slice. (B) TGIF coiled-coil prediction using Marcoils (left) and Ncols (right). The red line indicates predicted coiled-coil regions. (C) Schematic showing the TGIF genomic locus in parental and TGIF-Ty parasite lines. Primer binding sites used to confirm correct genomic integration are shown. (D) PCRs confirming Ty-AID integration into the TGIF genomic locus. (E) Schematic showing the Sec13 genomic locus in TGIF-Ty and TGIF-Ty:Sec13-GFP parasite lines made using CRISPR Cas9 editing tools. Position of gRNA and primer binding sites used to confirm correct genomic integration are shown. (F) PCRs confirming EmGFP integration into the Sec13 genomic locus

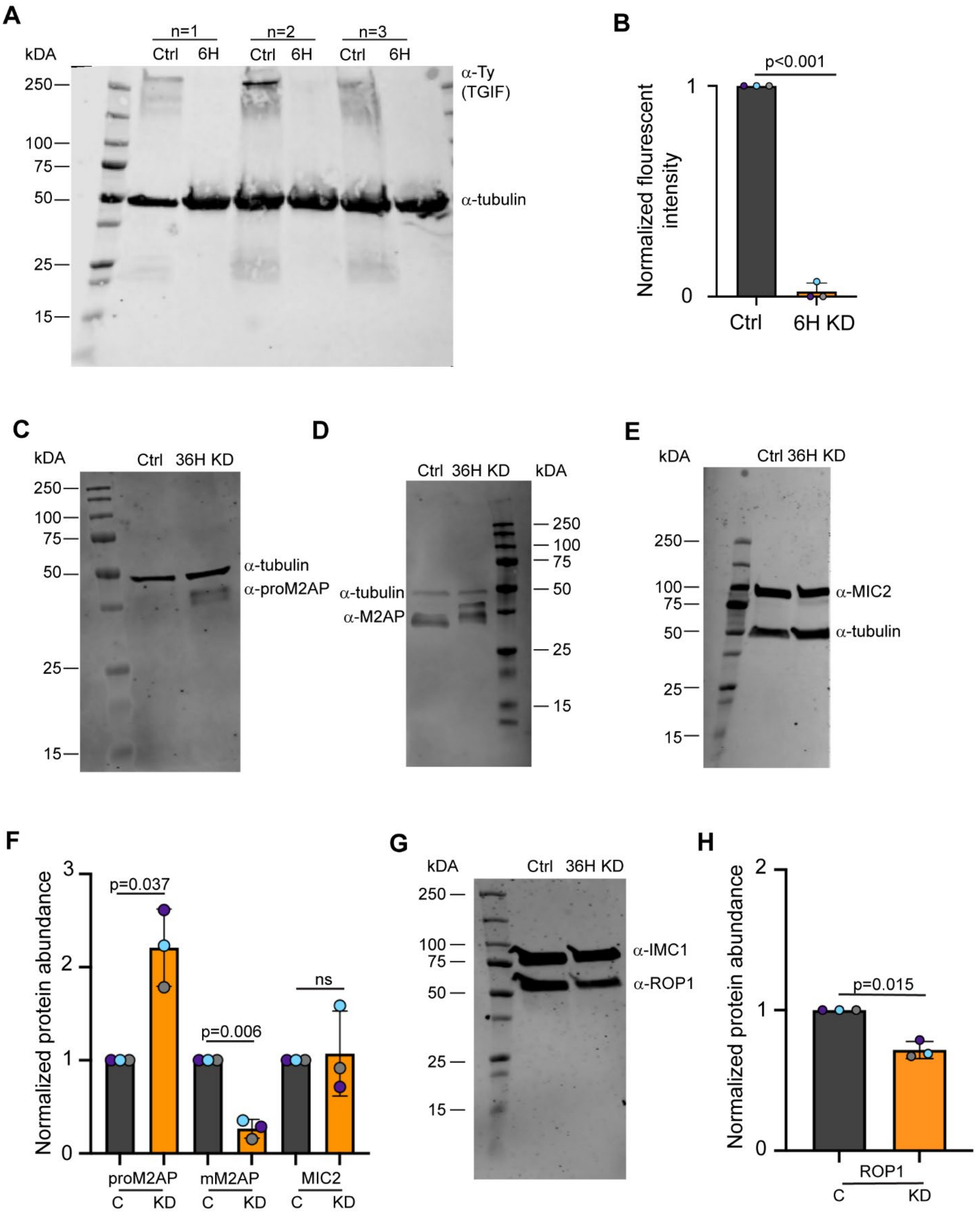

**Figure S2:** (A) Western blot on TGIF-Ty parasite lysates harvested after 6 hours of treatment with ethanol (control) or IAA (knockdown) in three experimental replicates. TGIF is 245kD and visualized using an anti-Ty antibody. Anti-tubulin (50kDa) is used as a loading control. (B) Quantification of western blot shown in (A). TGIF levels are depleted 99±0.02% after 6 hours of IAA treatment. (C-E) Western blot on TGIF-Ty parasite lysates harvested after 36 hours of treatment with ethanol (control) or IAA (knockdown) using anti-proM2AP (C), M2AP (D) and MIC2 (E) antibodies. Anti-tubulin (50kDa) is used as a loading control. (F) Quantification of proM2AP, M2AP and MIC2 protein levels in three independent replicates. (G) Western blot on TGIF-Ty parasite lysates harvested after 36 hours of treatment with ethanol (control) or IAA (knockdown) using an anti-ROP1 antibody. (H) Quantification of ROP1 protein levels in three independent replicates. In panels B, F and H statical significance was determined using a paired student's t=test. $P<0.05$ was considered significant.

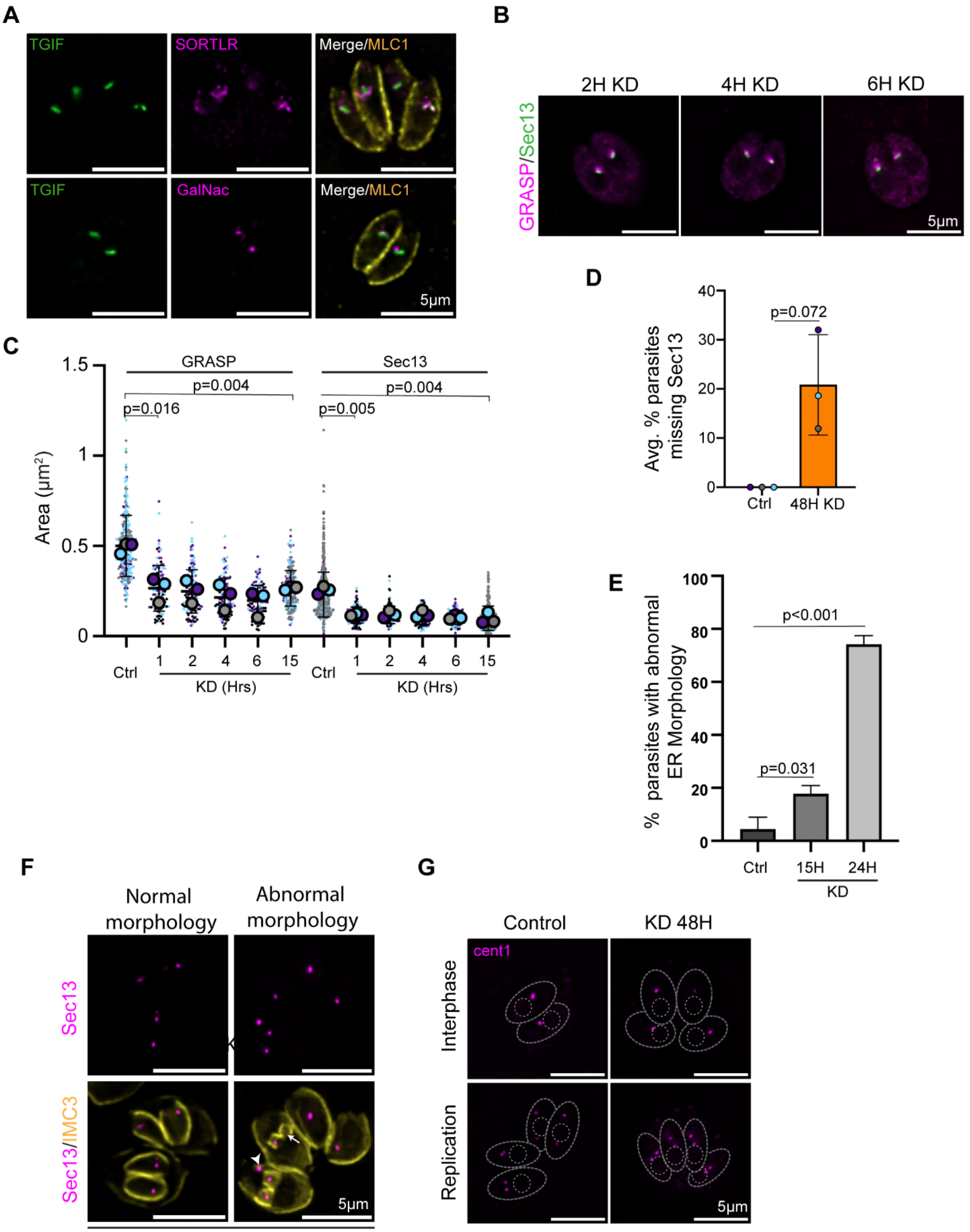

**Figure S3:** (A) Immunofluorescence of TGIF-Ty parasites. TGIF (anti-Ty: green) localizes adjacent to trans-golgi markers SORTLR (anti-SORTLR: magenta, top) and

GalNac-GFP (magenta, bottom). MLC1 (anti-MLC1) is shown in yellow. Scale bars are 5μm. Representative images from two experimental replicates (N = 20 vacuoles for each marker). (B) Immunofluorescence showing colocalization of mGRASP-mCherry (magenta) and Sec13-GFP (green) in TGIF-Ty:Sec13_GFP parasites treated with ethanol (control or IAA (knockdown) for 2, 4, and 6 hours. Scale bars are 5μm. Representative images from three experimental replicates (N > 35 vacuoles per condition). Images are maximum intensity projection of z-stacked images. (C) Quantification of Golgi area from images in Fig. 2A and Fig. S3B. For clarity, P-values from paired student t-test are indicated for 1- and 15-hour time points. Golgi area was significantly reduced at 2, 4 and 6 hour time points. (D) Bar chart showing average percent of parasites missing Sec13 after 48 hours of ethanol (control) or IAA (knockdown) treatment. Three experimental replicates were completed (N > 30 vacuoles per condition). P-values from paired t-tests are indicated. Colored circles data from each replicate. (E) Quantification of ER morphology from Fig. 2D. P-values from paired students t-tests are indicated. (F) Immunofluorescence showing Sec13-GFP (magenta) in TGIF-Ty:Sec13-GFP parasites treated with IAA (knockdown) for 48 hours. Anti-IMC3 (yellow) was used to visualize the periphery of daughter parasites. Parasites with normal and abnormal morphology are shown. White arrow indicates daughter missing Sec13. White arrowhead indicates Sec13 puncta outside daughter cells. Scale bar is 5μm. Representative images from three experimental replicates (N > 30 vacuoles per condition). (G) Immunofluorescence showing centrin1 fused with EmGFP (magenta) in TGIF-Ty parasites treated with ethanol (control) or IAA (knockdown) for 48 hours. Parasite outlines and nuclei are denoted by the grey dotted lines. Scale bars are 5μm. Representative images from two experimental replicates (N > 25 vacuoles per condition).

A

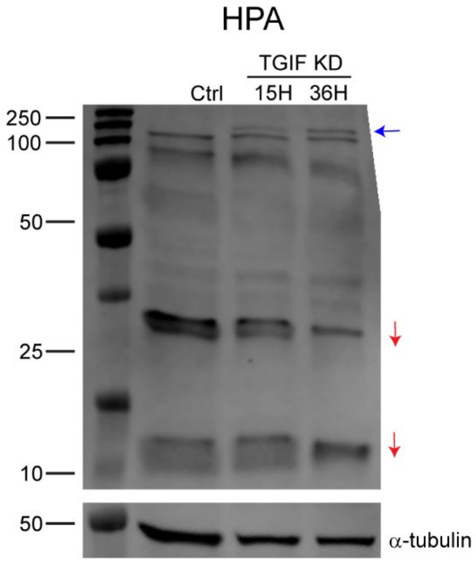

**Figure S4:** (A) Western blot with HPA, which recognizes O-glycans, and anti-tubulin (loading control) on TGIF-Ty parasite lysates harvested after 15 and 36 hours of IAA treatment, along with control parasites treated with ethanol. Additional bands are denoted with a blue arrow and loss of bands or decreased band intensity denoted with red arrows. Representative image from three experimental replicates.

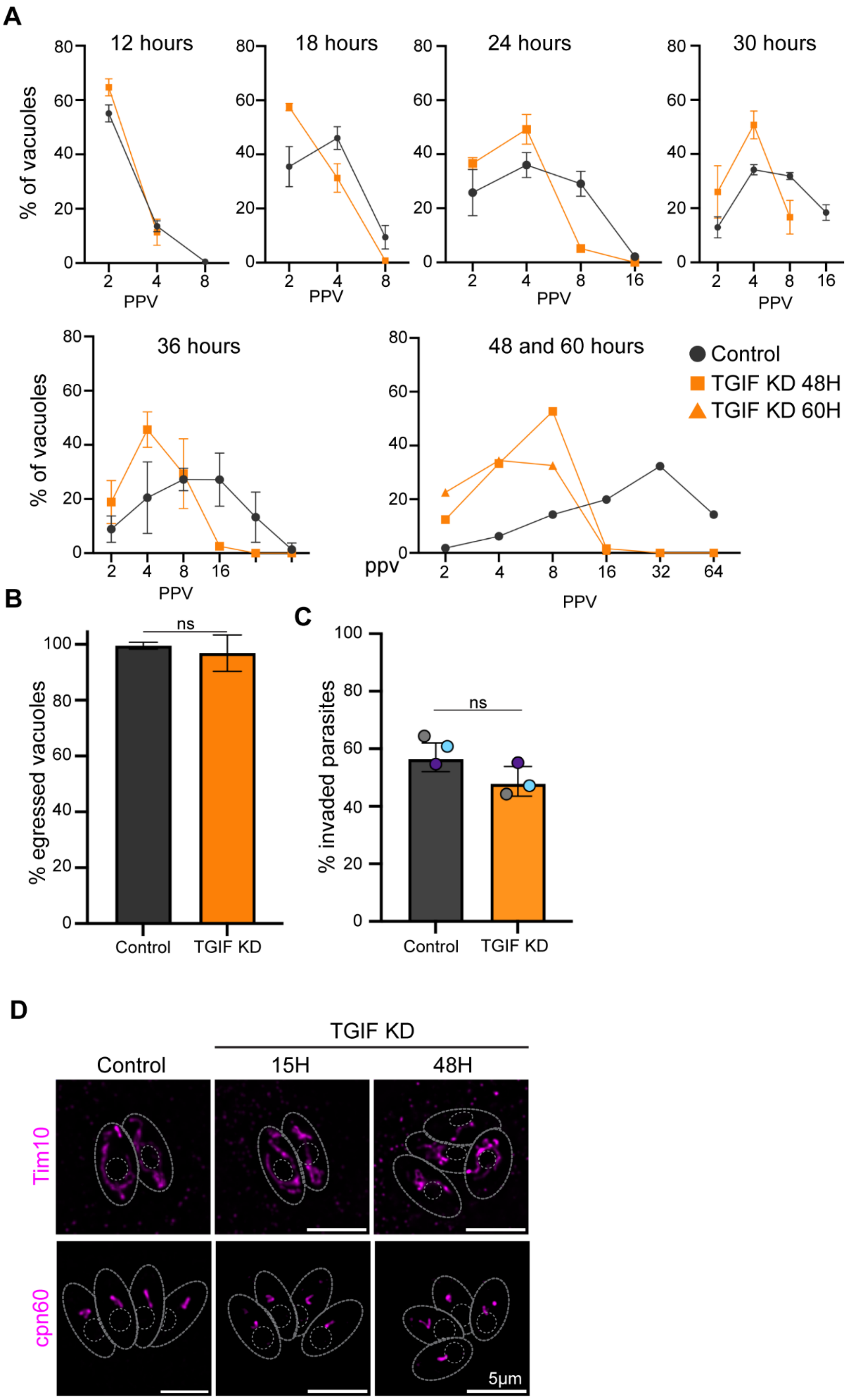

A
12 hours
18 hours
24 hours
30 hours
36 hours
48 and 60 hours
% of vacuoles
PPV
ppv
Control
TGIF KD 48H
TGIF KD 60H
B
% egressed vacuoles
ns
Control
TGIF KD
C
% invaded parasites
ns
Control
TGIF KD
D
TGIF KD
Control
15H
48H
Tim10
cpn60
5μm

**Figure S5:** (A) Graphs showing number of parasites per vacuole (PPV) after 12, 18, 24, 30, 36, 48 and 60 hours of growth in ethanol or IAA. Black lines and squares denote control parasites and orange lines and squares denote parasites treated with IAA. Orange lines and triangles denote parasites treated with IAA for 60 hours. (B) Bar chart showing percent of TGIF-Ty parasites that egressed from host cells upon treatment with A23187 after growth with ethanol (control) or IAA (knockdown) for 36 hours. 5 separate experiments were completed per condition for each experiment. (N > 200 vacuoles per condition). P-values from paired students t-tests are indicated. (C) Bar chart showing the percent of invaded TGIF-Ty parasites treated with ethanol (control) or IAA (knockdown) grown for 24 hours. (N = 4661 control parasites, and 3078 TGIF KD parasites were counted across three biological replicates). Colored circles indicate percent of parasites invaded in each replicate. P-values from paired students t-tests are indicated. (D) Immunofluorescence showing TGIF-Ty parasites expressing the mitochondrial marker Tim10-V5 visualized using anti-V5 antibody (magenta, top) and TGIF-Ty parasites stained with the apicoplast marker cpn60 (anti-cpn60, magenta, bottom) after treatment with ethanol (control) and IAA (knockdown) for 15 and 48 hours. Parasite outlines and nuclei are denoted by the grey dotted lines. Scale bars are 5μm. Representative images from two experimental replicates (N > 25 vacuoles per condition per marker).

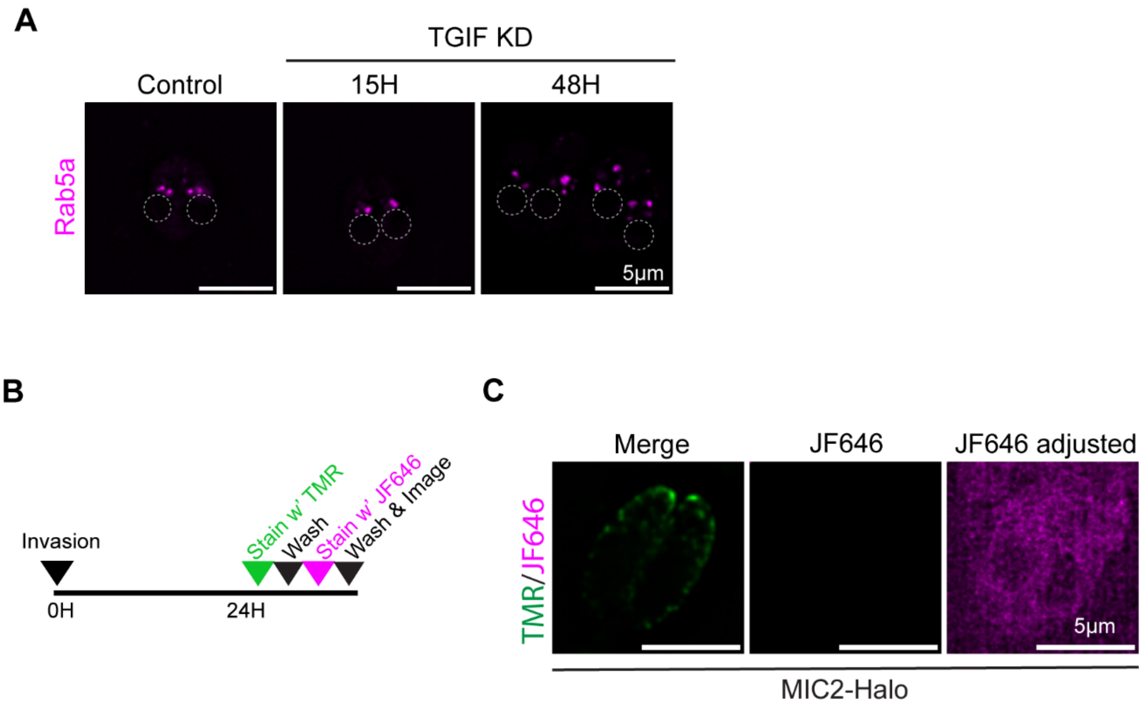

**Figure S6:** (A) Immunofluorescence showing the localization of Rab5a-NeonFP(magenta), a marker of the endosome-like compartment, in TGIF-Ty parasites grown with ethanol (control) or IAA (knockdown) for 15 and 48 hours. Parasite nuclei are indicated with a grey dotted circle. Scale bars are 5µm. Representative images from two experimental replicates (N = 20 vacuoles per condition).(B) Schematic of Halo labeling control experiment. (C) TGIF-Ty:Halo-MIC2 stained with HaloLigand-TMR (green) and Halo-Ligand-JF646 (magenta) as depicted in (B). Brightness and contrast of JF646 image is adjusted to show background staining. Representative images from one experimental replicate (N = 12 vacuoles).

**Supplementary Tables:**

| Gene (TgME49_) | PBS | Features | SP | TMD | Predicted Localization | CRISPR Score | References (PMID) |
|---|---|---|---|---|---|---|---|
| 224580 | A | RNA recognition motif-containing protein | no | no | nucleus/non-chromatin | -3.8 | |
| 224850 | A | polyadenylate-binding protein PABP | no | no | RNA granules | -4.12 | 29658303 |
| 228250 | B | elongation factor Tu GTP binding domain-containing protein | no | no | mitochondria | -3.39 | |
| 237015 | B | GRA43 | no | yes | dense granules/PVM | 2.4 | 38029261 |
| 266990 | C | beta-COP | no | no | ER/Golgi interface | -4.77 | 17485226 |
| 263200 | C | hypothetical protein | no | no | conoid/apical end | -0.61 | |
| **T213392** | **C** | **surface antigen repeat-containing protein** | **no** | **no** | **Nucleus-chromatin** | **-5.02** | |

**Table S1:** Hits from the Rab6 yeast-2-hybrid. Abbreviations: SP – signal peptide, TMD – transmembrane domain. PBS - predicted biological score.

| **Primer Name** | **Primer sequence** |
|---|---|
| | **TGIF-Ty** |
| TGIF gF1 | aaacgacggccagtgagcgcgcccaccgcggtggcctagggacgcgtttagactcgctct |
| TGIF gR1 | CGTCAAGTGGGTCCTGGTTAGTATGGACCTCCATagatctATTTCCATTTTCCCCCGCCC |
| F1 | gttcgcgtgggttctgcttc |
| R1 | ATTTCCATTTTCCCCCGCCC |
| R2 | AAGGGATCTTGATTCGTGTG |
| | **TGIF-Ty:Sec13-GFP** |
| Sec13_GFP protoF1 | AAGTTCCGATGTATGCGCCTTACAAg |
| Sec13_GFP protoR1 | aaaacTTGTAAGGCGCATACATCGGA |
| Sec13_GFP HR Oligo F | CGCGCCTCGGGCGCCGATGTATGCGCCTTACAAAGGAAACggactcgtgagcaagggcga |
| Sec13_GFP HR Oligo R (SG-R2) | agatttcttcgccttcctccttttcgctgaaaaggttTTActtgtacagctcgtccatgc |
| Sec13 PCR verification (SG-F1) | cattatgagcgtgcttctcc |
| Sec13 PCR verification (SG-R1) | GTTTCCTTTGTAAGGCGCAT |
| Sec13 guideRNA | CCGATGTATGCGCCTTACAA |
| | **TGIF-Ty:MIC2-Halo** |
| MIC2 HR Oligo F | AGAAGGAAACGCTGGTGCCCGTGGATGACGATAGTGATATGTGGATGGAGTCCGAAATCGGTACTGGCTT |
| MIC2 HR Oligo R | ctggtatgtgcgatctggaccagcgcaatctgctcaaagcgtcagtgctcTTAACCGGAAATCTCCAGAG |
| MIC2 guide RNA | TAGTGATATGTGGATGGAGT |
| | **TGIF-Ty:RON2-Halo** |
| RON2 HR Oligo F | TCAAGTACACGACCCCCGTCTTCCCCATGAGTGCGCCTCTCATCAAAGCCTCCGAAATCGGTACTGGCTT |

| | |
|---|---|
| RON2 HR Oligo R | gcgaaaagcgtccactgtctgtcctcctcgtccaacgccttcgtcttcttTTAACCGGAAATCTCCAGAG |
| RON2 guide RNA | aaacgtccgaaaaacagacg |
| **TGIF-Ty:ROP1-Halo** | |
| ROP1 HR Oligo F | AGGGCAGGCCCCTACTGGGCGAAGGAAGAGAGCAGGATGATGGATCGCAATCCGAAATCGGTACTGGCTT |
| ROP1 HR Oligo R | cgtttacgagaaacggtcttcgccgaatgcagcagcatgctgcccgtaTTAACCGGAAATCTCCAGAG |
| ROP1 guide RNA | TAAtacgggcagcatgctgc |

**Table S2:** List or primers used in this study.

| Plasmid name | FP | Localization |
|---|---|---|
| Ptub-GRASP-mCherry | mCherry | cis-Golgi |
| Ptub-SAG1-ΔGPI-GFP-HDEL | GFP | ER |
| Ptub-galnac-GFP | GFP | trans-Golgi |
| Pmin-centrin1-EmFP | EmGFP | centrosome |
| Pmin-EmFP-Rab6 | EmGFP | trans-Golgi |
| Pmin-Rab5a-neon | neonFP | PGCs |
| pmic8-Mic8-mCherry | mCherry | micronemes |
| Ptub-rop1-neon | neonFP | rhoptries |

**Table S3:** List of plasmids used in this study.

| Primary | | | | | | |
|---|---|---|---|---|---|---|
| **Target** | **Species** | **Concentration** | **Cellular location** | **Use** | **Reference (PMID)** | **Source** |
| IMC3 | mouse | 1:1000 | inner membrane complex | IFA | 15279956 | Dr. MJ Gubbels |
| MLC1 | rabbit | 1:1000 | inner membrane complex | IFA | | Yenzym: YZ8308/YZ8303 |
| SORTLR | mouse | 1:1000 | trans-Golgi | IFA | | Dr. Sabrina Marion |
| cent1 | mouse | 1:1000 | centrosome | IFA | | Kerafast: EBC004 |
| proM2AP | rabbit | 1:400/1:1000 | immature M2AP (microneme) | IFA/ WB | 16914527 | Dr. Vern Carruthers |
| M2AP | rabbit | 1:1000/ 1:10,000 | mature M2AP (microneme) | IFA/ WB | 11532123 | Dr. Vern Carruthers |
| MIC2 | rabbit | 1:500/1:5000 | microneme | IFA/ WB | 9208224 | Dr. Vern Carruthers |
| AMA1 | mouse | 1:1000 | microneme | IFA | 16000372 | Dr. Gary Ward |
| Cpn60 | rabbit | 1:1000 | apicoplast | IFA | 19808683 | Dr. Boris Striepen |
| ROP1 | mouse | 1:1000 | rhoptry bulb | IFA/ WB | | BEI resources: T52A3 |
| Ty | mouse | 1:1000 | Ty epitope | IFA/ WB | | Dr. de Graffenried |
| V5 | mouse | 1:1000 | V5 peptide | IFA | | chromotek: SV5-P-K |
| HPA | n/a | 1:250 | O-glycosylated proteins | WB | 17652409 | ThermoFisher: L32454 |
| Tubulin | mouse | 1:5000 | tubulin | WB | | Sigma T6074 |
| SAG1 | mouse | 1:100 | Plasma membrane | IFA | | abcam: 156836 |
| IMC1 | mouse | 1,1000 | inner membrane complex | WB/I FA | | Dr. Gary Ward |

| | | | | | | |
|---|---|---|---|---|---|---|
| **Secondary** | | | | | | |
| **Target** | **Species** | **Concentration** | **Fluorophore** | | | |
| Anti-mouse | goat | 1:1000 | AlexaFluor-488 | IFA | | Invitrogen: A11001 |
| Anti-mouse | goat | 1:1000 | AlexaFluor-546 | IFA | | Invitrogen: A11030 |
| Anti-mouse | goat | 1:1000 | AlexaFluor-647 | IFA | | Invitrogen: A21235 |
| Anti-rabbit | goat | 1:1000 | AlexaFluor-488 | IFA | | Invitrogen: A11034 |
| Anti-rabbit | goat | 1:1000 | AlexaFluor-546 | IFA | | Invitrogen: A11035 |
| Anti-rabbit | goat | 1:1000 | AlexaFluor-647 | IFA | | Invitrogen: A21245 |
| Anti-mouse | goat | 1:1000 | IR Dye 680 | WB | | Licor: 926-68070 |
| Anti-rabbit | goat | 1:1000 | IR Dye 800 | WB | | Licor: 926-32211 |

**Table S4:** List of antibodies used in this study.

**Supplementary Video Legends:**

**Video S1: Egress assay with control parasites.** TGIF-Ty parasites grown with ethanol (control) for 36 hours before being treated with calcium ionophore to induce egress. DIC images were taken every second for 5 minutes. 7 separate experiments were completed (N = 204 vacuoles). Scale bars is 10μM. Time in seconds after addition of ionophore is indicated.

**Video S2: Egress assay with TGIF knockdown parasites.** TGIF-Ty parasites grown with IAA (TGIF KD) for 36 before being treated with calcium ionophore to induce egress. DIC images were taken every second for 5 minutes. 10 separate experiments were completed. (N = 273 vacuoles). Scale bars is 10μM. Time in seconds after addition of ionophore is indicated.
